# NuMA interaction with chromatin is vital for proper nuclear architecture in human cells

**DOI:** 10.1101/2020.05.02.073767

**Authors:** Ashwathi Rajeevan, Riya Keshri, Sachin Kotak

## Abstract

NuMA is an abundant long-coiled-coil protein that plays a prominent role in spindle organization during mitosis. In interphase, NuMA is localized to the nucleus and hypothesized to control gene expression and chromatin organization. However, because of the prominent mitotic phenotype upon NuMA loss, its precise function in the interphase nucleus remains elusive. Here, we report that NuMA is associated with chromatin in interphase and prophase but released upon nuclear envelope breakdown (NEBD) by the action of Cdk1. We uncovered that NuMA directly interacts with DNA via evolutionarily conserved sequences in its C-terminus. Notably, the expression of the DNA-binding mutant of NuMA affects chromatin decondensation at the mitotic exit, and nuclear shape in interphase. The impact on nuclear shape by mutant NuMA expression is due to its potential to polymerize into high-order fibrillar structures. This study links the chromatin binding ability of NuMA with the maintenance of nuclear shape and architecture, which has a well-studied role in regulating gene expression during development and diseases.

## Introduction

In a eukaryotic cell, the nucleus is the largest organelle that harbors the genetic information and non-membrane organelles that are essential for the existence of life. Recent work has shown that the proper structural organization and the mechanical properties of the nucleus are vital for ensuring error-free functioning of various cellular processes by directly controlling proper gene regulation (Lammerding et al., 2004; Finn et al., 2019; Nagano et al., 2017; Nozaki et al., 2017; reviewed in Friedl et al., 2011; Van Steensel and Belmont, 2017; Mirny et al., 2019). Within the nucleus, chromatin is organized in a non-random manner into defined regions called chromosomal territories. Any perturbation in the nuclear architecture can influence these territories, which in turn can impact gene expression and the cell cycle (Smith et al., 2017; Aureille et al., 2019; reviewed in Webster et al., 2009; Cremer and Cremer, 2010). Moreover, these territories must also be preserved while chromatin undergoes condensation and decondensation cycle during mitosis (reviewed in Antonin and Neumann, 2016). Despite the identification of a few proteins that maintain the proper nuclear architecture and ensures correct chromatin states during mitosis, our thorough understanding in this remarkable feat is far from complete (reviewed in Misteli, 2007; Simon and Wilson, 2011; Hubner et al., 2013; Antonin and Neumann, 2016).

The Nuclear Mitotic Apparatus (NuMA) is a large protein (2115 amino acids) with two globular domains separated by a long coiled-coil domain (Yang et al., 1992). It is estimated that approximately 10^6^ molecules of NuMA are present in mammalian cells (Compton et al., 1992; reviewed in Cleveland, 1995). NuMA is present in the nucleus during interphase. However, upon nuclear envelope breakdown (NEBD) in mitosis, it localizes at the spindle poles and cell cortex, where it is required for the proper assembly and maintenance of the mitotic spindle as well as spindle orientation and elongation (Lydersen and Pettijohn, 1980; Yang and Snyder, 1992; Compton and Cleveland, 1993; Merdes et al., 1996; Merdes et al., 2000; Woodard et al., 2010; Kiyomitsu and Cheeseman, 2012; Kotak et al., 2012; Seldin et al., 2013; Kotak et al., 2014; Zheng et al., 2014; Hueschen et al., 2019). Because NuMA present as an abundant protein in interphase nuclei, several research groups have studied NuMA’s function in the nucleus (reviewed in Radulescu and Cleveland, 2010). Within the nucleus, NuMA was proposed to be a part of a nuclear matrix, i.e., insoluble 3D-network resistant to nucleases and high-salt (Price and Pettijohn, 1986; Zeng et al., 1994; Merdes and Cleveland, 1998). This finding was further supported by structural studies showing that NuMA can form multiarm oligomers with its coiled-coil and C-terminal domain, and overexpression of NuMA creates a quasi-hexagonal organization that can fill the nuclei (Harborth et al., 1999). Additionally, NuMA has been shown to co-localize with several nuclear proteins, including high mobility group proteins (HMG I/Y), transcription factor GAS41 and p53, suggesting its role in gene regulation (Harborth et al., 2000; Tabellini et al., 2001; Endo et al., 2013). In this realm, microinjection of anti-NuMA antibodies or expression of the truncated form of NuMA caused nuclear shape and organization defects (Kallajoki et al., 1991; Kallajoki et al., 1993; Compton and Cleveland, 1993; Gueth-Hallonet et al., 1998). However, whether these defects were due to compromised mitosis upon NuMA inactivation remained unknown (reviewed in Radulescu and Cleveland, 2010). Also, there is no clear evidence that NuMA interacts with chromatin inside the nucleus of a living cell. Even, if it does, what is the biological significance of this interaction, and how it gets released from chromatin upon NEBD remained unexplored.

In this study, we show that NuMA is associated with chromatin inside the nucleus during interphase. At the mitotic onset, Cdk1/cyclinB1 (referred to as Cdk1)-mediated phosphorylation in late-prophase releases NuMA from chromatin. Importantly, we identify an evolutionarily conserved domain rich in arginine and lysine residues at its C-terminus that is responsible for NuMA-DNA interaction. Moreover, the expression of a mutant NuMA lacking the DNA binding potential impact chromatin decompaction during nuclear envelope reformation (NER). Additionally, this mutated NuMA undergoes higher-order assemblies and forms puncta and solid fibrillar structure that perturbs nuclear shape. Overall, this study uncovers a novel role of NuMA in maintaining the proper nuclear architecture that is independent of its mitotic function.

## Results and Discussion

### NuMA interacts with chromatin in the interphase nucleus

To investigate the mobility of NuMA in the interphase nucleus, we sought to conduct fluorescence recovery after photobleaching (FRAP) analysis in the HeLa Kyoto cell line that stably expresses AcGFP (*Aequora coerulescens* GFP), and a mono-FLAG tagged NuMA (AcGFP-NuMA; Fig. 1A). This engineered line expresses the AcGFP-NuMA amount that was comparable to that of the endogenous protein (Fig. 1B). siRNAs-mediated depletion of endogenous NuMA led to chromosome instability and the appearance of chromosome bridges in a significant number of cells during mitosis, and these phenotypes were completely suppressed in AcGFP-NuMA line, indicating that this cell line is operational (Supplementary Fig. S1A and S1B). FRAP analysis in this line revealed that the half-time for the recovery [t_1/2_] of AcGFP-NuMA is about ∼13 s (Fig. 1C, 1F, and 1I). AcGFP-NuMA [t_1/2_] values are in stark contrast in comparison with robustly diffusing AcGFP-NLS or tightly associated histone protein AcGFP-H2B where the [t_1/2_] values were estimated ∼1.5 s and 29 s, respectively (Fig. 1I; Supplementary Fig. S1C-S1F). Notably, [t_1/2_] of AcGFP-NuMA is analogous to various transcription factors that are transiently associated with the DNA (Sekiya et al., 2009; reviewed in Houtsmuller, 2005; Mueller et al., 2010).

**Figure 1.**
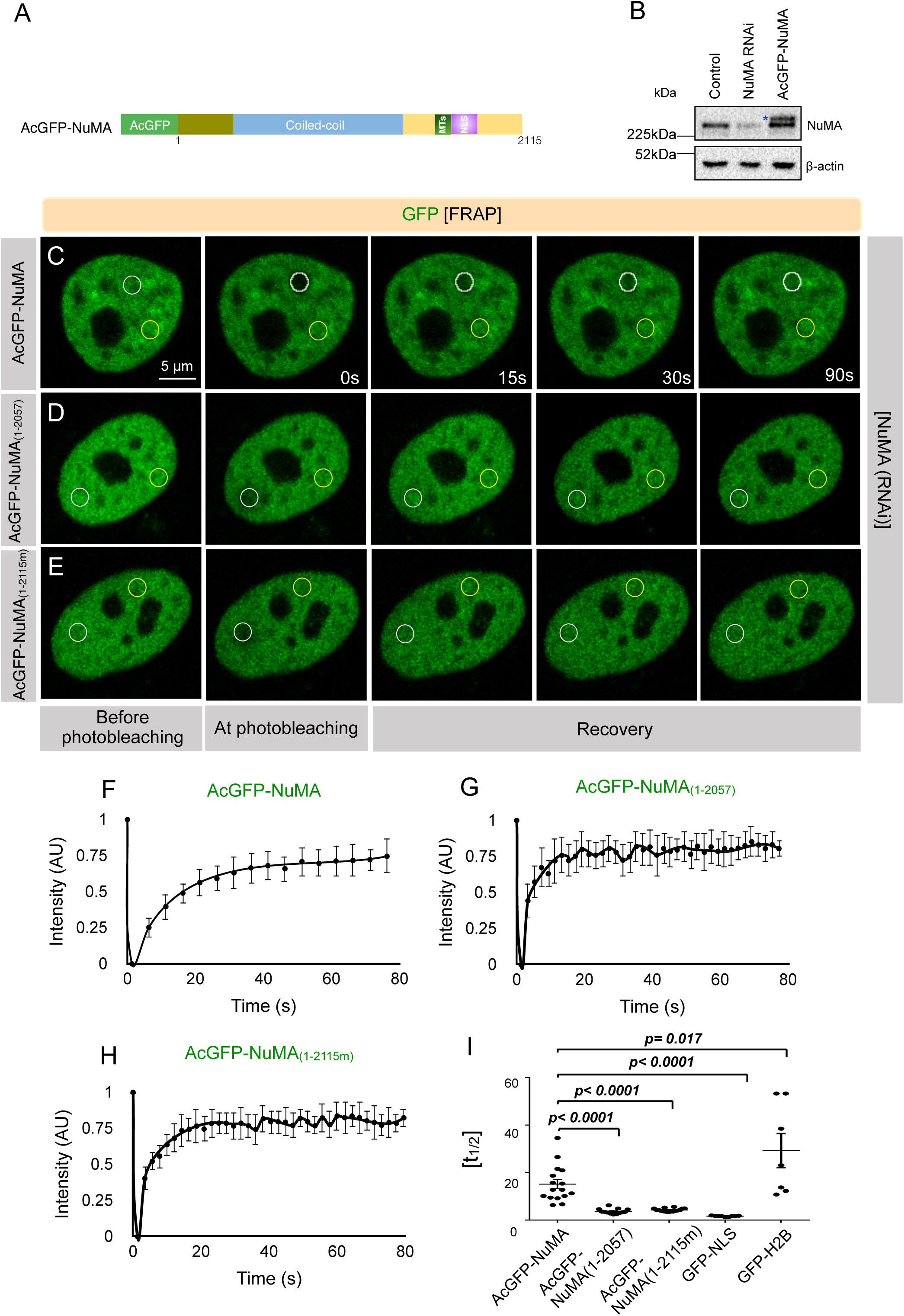
NuMA is transiently associated with chromatin in the interphase nucleus. (A) Domain organization of NuMA with mono FLAG (FL) and AcGFP-tag at the N-terminus (referred to as AcGFP-NuMA). The coiled-coil domain, the region mediating interaction with microtubules (MTs), and the nuclear localization signal (NLS) are depicted. (B) Immunoblot analysis of protein extracts prepared from the mitotically synchronized HeLa Kyoto cells, which are transfected with scrambled siRNAs (Control), siRNAs against NuMA 3’-UTR for 72 hr, or left untreated and stably expressing AcGFP-NuMA. Extracts were probed with antibodies against NuMA and β-actin. Transgenic AcGFP-NuMA protein is shown by a blue asterisk that is migrating above the endogenous protein. The molecular mass is indicated in kilodaltons (kDa). (C-I) FRAP analysis of HeLa Kyoto cells that are stably expressing AcGFP-NuMA (C, F), transiently transfected with AcGFP-NuMA_(1-2057)_ (D, G) or AcGFP-NuMA_(1-2115m)_ (E, H) and are depleted for endogenous NuMA. The GFP signal is shown in green, and the time is indicated in seconds (s). The unbleached and bleached region of the cell is shown by yellow and white circles, respectively. The GFP recovery profile of the bleached area corrected for photobleaching is plotted for 80s for all three conditions. Note the half-time of recovery [t_1/2_] of cells expressing AcGFP-NuMA is ∼13s, which is remarkably slow in comparison with AcGFP-tagged NLS (nuclear localization signal) expressing cells [t_1/2_=∼1.5s], and significantly faster in contrast to AcGFP-H2B expressing cells [t_1/2_=∼29s] (I). Analogous [t_1/2_] value (∼12.2s) was obtained in cells which are transiently transfected with AcGFP-NuMA (data not shown). Also, note that rapid GFP recovery in AcGFP-NuMA_(1-2057)_ [t_1/2_=∼3.6s] and AcGFP-NuMA_(1-2115m)_ [t_1/2_=∼4.2s] in comparison to that of AcGFP-NuMA expressing cells (I). Statistical significance is calculated by unpaired t-test. (n>10 for all the cases except AcGFP-H2B expressing cells where n=7; Error bars: SD for F-H and SEM for I)

To corroborate this finding with a biochemical method, we isolated chromatin and nuclear matrix (non-chromatin, ribonucleoproteinaceous framework that is resistant to high salt) from interphase nuclei and we analyzed the association of NuMA in these fractions. As reported earlier, we uncovered that NuMA is associated with the nuclear matrix in HeLa (Supplementary Fig. S1G; Zeng et al., 1994; Das et al., 1993; Abad et al., 2007). Interestingly, a significant portion of NuMA was also associated with chromatin fraction, as reported for mammary epithelium cells [Supplementary Fig. S1G; Abad et al., 2007]. Altogether these data suggest that NuMA is not freely diffusing inside the interphase nuclei.

### Cdk1-mediated phosphorylation release NuMA from chromatin at mitotic entry

During interphase, NuMA is present in the entire nucleoplasm, except for nucleoli (Lydersen and Pettijohn, 1980; Kallajoki et al., 1992; Tang et al., 1993). However, in mitosis, NuMA is restricted to the spindle poles and the cell cortex (Lydersen and Pettijohn, 1980; Compton et al., 1992; Compton and Cleveland, 1993; Merdes et al., 1996; Du and Macara, 2004; Woodard et al., 2010; Kiyomitsu and Cheeseman, 2012; Kotak et al., 2012). Data obtained from FRAP and biochemical analysis told us that NuMA might associate with chromatin in interphase. Thus, we wondered if we can visualize this interaction in a living cell. Chromosomes are present as a contiguous mass in the interphase nuclei and appear as distinct bodies during late prophase (reviewed in Batty and Gerlich, 2019). Therefore, we sought to examine the localization of NuMA in synchronized non-transformed hTERT-RPE1 cells during late prophase before NEBD (Fig. 2A). Importantly, we found that NuMA significantly enriches onto the chromosomes in late prophase, and colocalizes with another chromatin-associated protein RanGEF RCC1 (Supplementary Fig. S2A). Similar results were obtained in HeLa (data not shown). To scrutinize this further, we performed a super-resolution analysis of NuMA, DNA, and RCC1 in the late prophase cells. This data showed that NuMA enriches at the periphery of condensed chromatin in late prophase cells and colocalizes with RCC1 (Fig. 2B). Importantly, we also uncovered that the GFP-tagged C-terminus of NuMA [GFP-NuMA_(1411-2115)_] localizes to chromatin in late prophase similar to endogenous proteins, suggesting NuMA interacts with chromatin through its C-terminus (Supplementary Fig. S2D).

**Figure 2.**
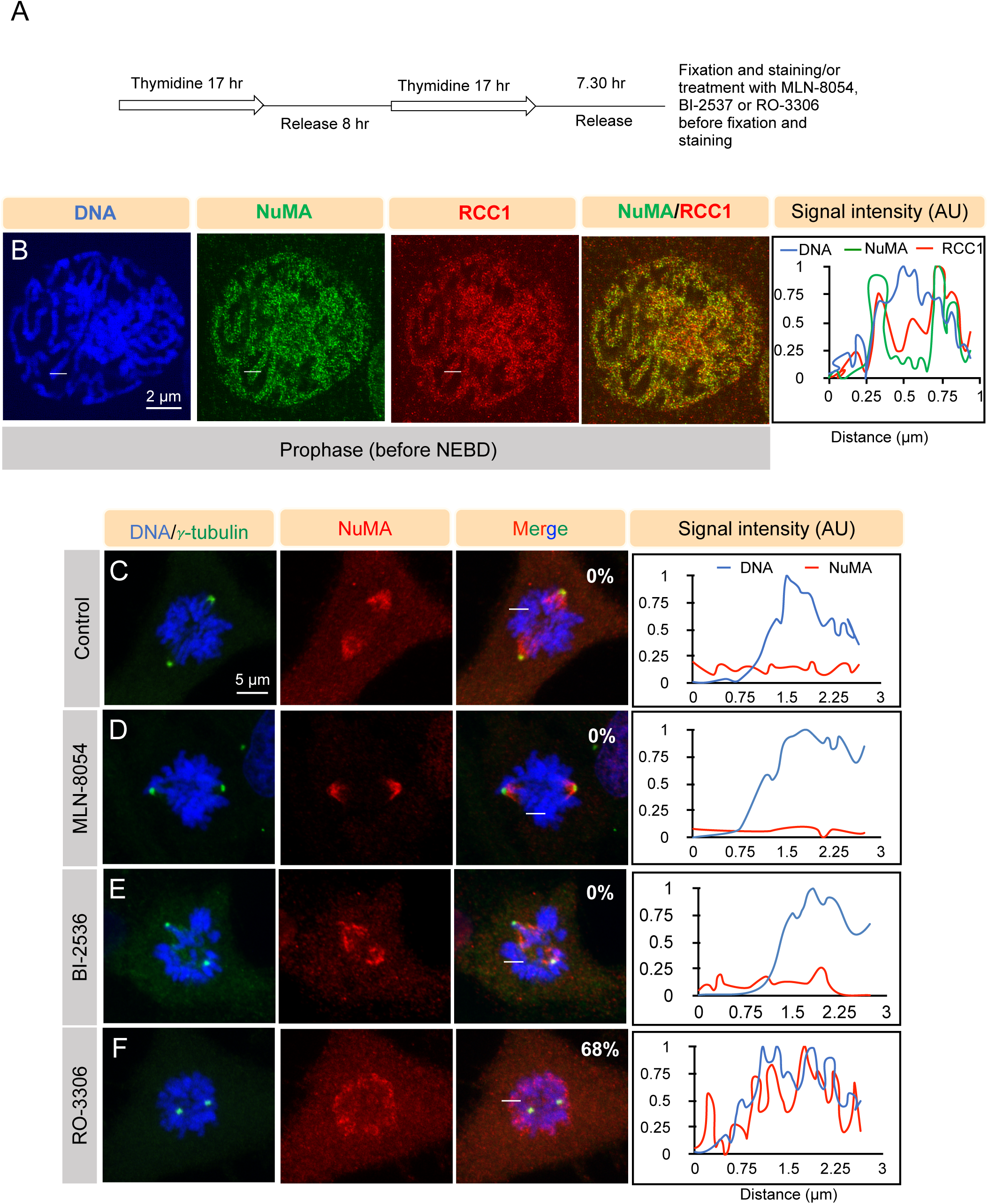
Cdk1 activity is critical for releasing NuMA from chromatin upon mitotic entry. (A) hTERT-RPE1 cell synchronization scheme for enriching cells in the late prophase following double thymidine release. Cells were fixed after 7.30 hr of double thymidine release for obtaining maximum number of cells in prophase. Cells were treated with DMSO (Control), Aurora A inhibitor MLN-8054 (250 nM for 1 hr), Plk1 inhibitor BI-2536 (300 nM for 30 min) or Cdk1 inhibitor RO-3306 (20 µM for 10 min) before fixation. (B) Super-resolution images of prophase synchronized hTERT-RPE1 cells immunostained for NuMA (green) and RanGEF RCC1 (red). DNA is visualized in blue. In this and other Figures, a white line on the confocal images represent the area that is utilized to make a line scan plot shown on the right. (C-F) hTERT-RPE1 cells synchronized in late prophase, as indicated in panel A, are treated either with DMSO control (C), MLN-8054 (D), BI-2536 (E) or RO-3306 (F). After fixation, these cells are stained for NuMA (red) and γ-tubulin (green). The percentage of cells showing chromosomal retention of NuMA in cells treated with the various inhibitor is indicated on the corresponding images. Note the retention of NuMA on chromatin in prometaphase cells that were treated with Cdk1 inhibitor RO-3306 compared to the control cells. Also, check Supplementary Figure S3 for control experiments. (n>20 cells in each condition and experiments were repeated four times).

By utilizing a temperature-sensitive hamster cell line tsBN2 that is affected for RCC1 (Nishimoto et al., 1978; Nishitani et al., 1991), it was hypothesized that NuMA might interact with RCC1 or RCC1-dependent protein (Compton and Cleveland, 1993). Since NuMA shows significant co-localization with RCC1, we investigated if NuMA or RCC1 are interdependent for their chromatin localization in late prophase nuclei.

Interestingly, RNAi-mediated depletion of RCC1 or NuMA did not perturb NuMA or RCC1 localization onto chromatin, indicating that NuMA localization on chromatin is independent of RanGEF RCC1 (compare Supplementary Fig. S2B and S2C with S2A). During mitotic entry, phosphorylation by several mitotic kinases is critical for remodeling the mitotic proteome and, thereby, the localization of a number of proteins (reviewed in Lindqvist et al., 2009; Cuijpers and Vertegaal, 2018). Thus, we decided to identify the kinase that is responsible for releasing NuMA from chromatin upon NEBD with high temporal resolution. To this end, we inactivated Aurora A kinase, Polo-like kinase (Plk1), and Cdk1 in a synchronized G2/prophase population of hTERT-RPE1 cells using MLN-8054, BI-2536 and RO-3306 respectively, and analyzed NuMA localization after NEBD in prometaphase (Figure 2A; Vassilev et al., 2006; Hoar et al., 2007; Steegmaier et al., 2007). Interestingly, inactivation of Cdk1, but not Aurora A or Plk1 caused the retention of NuMA on the chromosomes during prometaphase (Fig. 2C-2F). The inability of Aurora A or Plk1 inhibition to restore NuMA localization cannot be due to partial inactivation of these kinases as we noted a robust impact of MLN-8054 and BI-2536 on spindle pole localization of NuMA and central spindle localization of a RhoGEF ECT2 as reported previously (Supplemental Fig. S3A-S3F; Petronczki et al., 2007; Kotak et al., 2016). And, identical results were obtained with a fivefold higher dose of these inhibitors (Supplementary Fig. S3G-S3J) Next, to test if direct phosphorylation of NuMA by Cdk1 would release NuMA from chromatin, we mutated nine threonine or serine residues which were identified in recent phosphoproteomics data set to alanine (Supplementary Fig. S3K) (Dephoure et al., 2008; Petrone et al., 2016; Rogers et al., 2015). However, we failed in identifying a single Cdk1 site that uncouples NuMA from chromatin (please see discussion).

Whereas the mechanism whereby Cdk1 phosphorylation uncouples NuMA from the chromatin upon mitotic entry will be of interest for future work, our data support the notion that Cdk1 activity is vital for releasing NuMA from the chromatin upon mitotic entry.

### NuMA directly interacts with the DNA through its C-terminus

Analogous to the endogenous protein, GFP-NuMA_(1411-2115)_ localizes on the chromatin in prophase but not during prometaphase and metaphase (Fig. 3A and 3B; Supplementary Fig. S2D and S4A; Supplementary Movie S1; Kotak et al., 2013; Sana et al., 2018).

**Figure 3.**
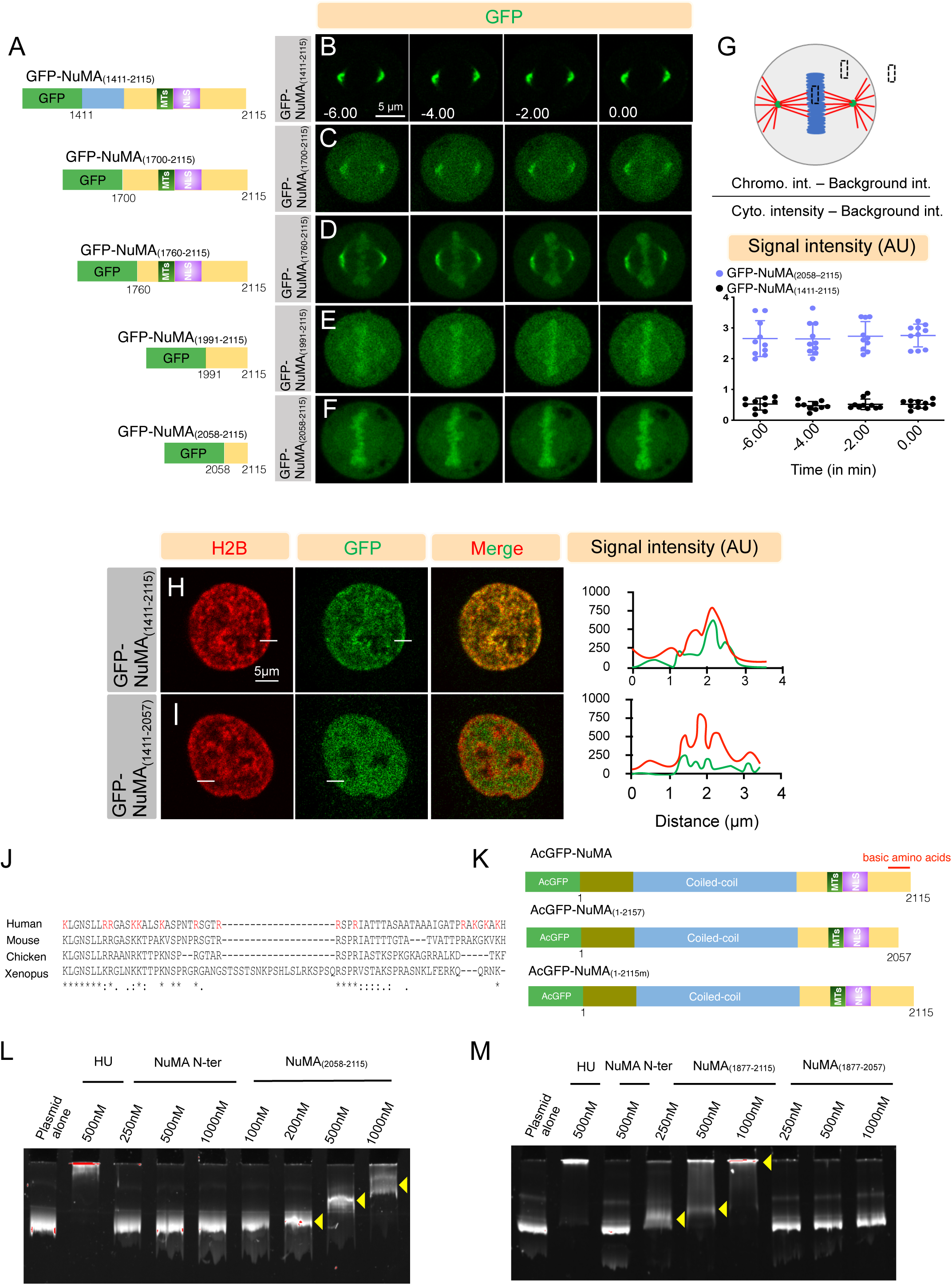
NuMA interacts with the DNA with the evolutionarily conserved region present in its C-terminus. (A) Schematic representation of GFP-tagged NuMA constructs used for the experiments that are shown on the right; the regions mediating interaction with microtubules (MTs), and the nuclear localization signal (NLS) are represented. (B-F) Images from time-lapse recording of HeLa cells stably expressing mCherry-H2B and transiently transfected with GFP-NuMA_(1411-2115)_ (B), GFP-NuMA_(1700-2115)_ (C), GFP-NuMA_(1760-2115)_ (D), GFP-NuMA_(1991-2115)_ (E) or GFP-NuMA_(2058-2115)_ (F). The GFP signal is shown in green. Time is indicated in minutes with t=0 corresponding to the last frame of metaphase before the onset of chromosomes segregation. Note the enrichment of GFP signal on the metaphase chromosome for cells expressing GFP-NuMA_(1760-2115)_, GFP-NuMA_(1991-2115)_, and GFP-NuMA_(2058-2115)_. (G) Chromosomal intensity quantification scheme of a metaphase cell; black boxes indicate the area used for the quantification of the signal intensity. The ratio of the chromosomal to cytoplasmic GFP-signal intensity is plotted over time for GFP-NuMA_(1411-2115)_ and GFP-NuMA_(2058-2115)_. p<0.0001 between GFP-NuMA_(1411-2115)_ and GFP-NuMA_(2058-2115)_ for all the time points studied. Statistical significance is calculated by unpaired-t-test. (n=10 cells for all cases; Error bars: SD). (H, I) Images from the time-lapse recording of HeLa cells in prophase before nuclear envelope breakdown (NEBD) that are stably expressing mCherry-H2B and transiently transfected with GFP-NuMA_(1411-2115)_ (H), or GFP-NuMA_(1411-2057)_ (I). Note that the GFP signal is homogeneously distributed in the nucleus in GFP-NuMA_(1411-2057)_ expressing cells in comparison to the cells expressing GFP-NuMA_(1411-2115)_ where the signal is localized to chromatin. Linescan plot is shown on the right. (J) Sequence alignments of NuMA DNA binding region (2058-2115) with NuMA orthologs. (*Homo sapiens* NM_006185.2, *Mus musculus* NP_598708.3, *Gallus gallus* NP_001177854.1, *Xenopus laevis* NP_001081559.1.) The basic amino acids (Arginine and Lysine residues) are highlighted in red. Note that majority of basic amino acids are conserved across these species. (K) Schematic representation of AcGFP-tagged full-length NuMA (AcGFP-NuMA) or NuMA that is either deleted [AcGFP-NuMA_(1-2057)_] or mutated [AcGFP-NuMA_(1-2115m)_] for the last 58 aa. The coiled-coil domain, the region mediating interaction with microtubules (MTs), and the nuclear localization signal (NLS) are shown. (L, M) Gel mobility shift assay of pUC19 plasmid (400ng) that is incubated with the indicated concentration of *E. coli* generated recombinant proteins against bacterial histone-like protein HU, Hexa-histidine-NuMA N-ter (indicated as NuMA N-ter), and Hexa histidine-NuMA_(2058-2115)_ [indicated as NuMA_(2058-2115)_] (L). Or with HU, NuMA N-ter, Hexa histidine-NuMA_(1877-2115)_ [indicated as NuMA_(1877-2115)_], and Hexa histidine-NuMA_(1877-2057)_ [(indicated as NuMA_(1877-2057)_] (M). Yellow arrowheads indicate the retardation of pUC19 plasmid DNA upon the increasing concentration of NuMA_(2058-2115)_ and NuMA_(1877-2115)_, but not with NuMA_(1877-2057)_ missing the last 58 aa.

Therefore, we decided to identify a minimum signature sequence of NuMA in its C-ter that would enable it to associate with the chromatin independent of the mitotic stages. To this end, we generated several GFP-fusion C-ter fragments of NuMA and analyzed their localization at various stages of mitosis (Fig. 3A). Interestingly, in contrast to NuMA_(1411-2115)_, the expression of NuMA_(1760-2115)_ and NuMA_(1991-2115)_ showed significant enrichment on chromatin even in prometaphase and metaphase stages (compare Fig. 3D and 3E with 3B; Supplementary Fig. S4A-S4D). This analysis further demonstrated that the sequence comprising of the last 58 amino acids (2058-2115) is both necessary and sufficient for its interaction with the chromatin in prophase as well as in metaphase (Fig. 3F-3I; Supplementary Fig. S4E: Supplementary Movies S2 and S3). Similarly, the robust localization of NuMA_(1760-2115)_ on metaphase chromosomes is lost upon the deletion of the last 58 amino acids (Supplementary Fig. S4F-S4H). Altogether, these data indicate that the ability of NuMA to interact with chromatin is due to the presence of the last 58 aa in its C-ter.

Sequence 2058-2115 is evolutionarily conserved, and rich in the positively charged arginine and lysine residues (Fig. 3J). Thus, one possibility would be that these amino acids directly interact with acidic DNA sequences, enabling NuMA-DNA interaction. Notably, *Escherichia coli* generated Hexa-histidine-tagged 58 aa recombinant protein [6HIS-NuMA_(2058-2115)_], but not a recombinant protein comprising of either NuMA N-terminus, or C-ter lacking the last 58 aa could interact with plasmid DNA in a gel mobility shift assay (Fig. 3L and 3M).

Further, to scrutinize the role of NuMA’s DNA binding ability in the context of the full-length protein and the biological significance of this interaction, we conducted FRAP analysis with the AcGFP-tagged full-length NuMA lacking the last 58 aa [AcGFP-NuMA_(1-2057)_; Fig. 3K]. Remarkably, FRAP analysis revealed that AcGFP-NuMA_(1-2057)_ shows greater mobility in the interphase nucleus when compared with AcGFP-NuMA full-length protein (Fig. 1D, IG, and 1I). This observation suggests that the last 58 aa are crucial for their interaction with the chromatin, and possibly this interaction restricts NuMA mobility inside interphase nuclei. Moreover, FRAP analysis with a mutant NuMA where all the fourteen basic amino acid residues present between 2058-2115 are mutated to alanine [AcGFP-NuMA_(1-2115m)_; Fig. 3K] showed dramatic mobility of the mutated NuMA construct in comparison to the wild-type protein (Fig. 1E, IH, and 1I).

### NuMA-DNA interaction is critical for proper chromosomes decondensation at the mitotic exit

Upon nuclear envelope reformation (NER), NuMA localizes back to the nucleus because of the presence of nuclear localization signal (NLS) in the C-ter (reviewed in Cleveland, 1995; Radulescu and Cleveland, 2010). Because AcGFP-NuMA_(1-2057)_ lacks the DNA binding motif, we sought to analyze the relevance of NuMA-chromatin interaction upon NER. Importantly, in comparison with the full-length protein, expression of either AcGFP-NuMA_(1-2057)_ or AcGFP-NuMA_(1-2115m)_ in NuMA (RNAi) background did not lead to sufficient chromosomes decondensation in late mitosis (Fig. 4A-4D; Supplementary Movies S4-S6). This impact on the chromosomes decompaction in cells expressing either the deleted or mutated version of the protein is not because of its inability to localize to the nucleus in comparison to the wild-type protein (data not shown). To further evaluate the importance of NuMA’s DNA binding ability for chromosomes decompaction and its impact on the nuclear shape in the early G1 phase of the cell cycle, we imaged HeLa cells that are stably expressing nuclear envelope marker mCherry-LaminB1 and are transiently transfected either with the wild-type or AcGFP-NuMA_(1-2057)_ in NuMA (RNAi) background after anaphase onset (Fig. 4E and 4F). Interestingly, the nuclear width of the cells expressing NuMA that lacks DNA binding ability is significantly smaller in comparison with the cells that express the wild type protein (Fig. 4G).

**Figure 4.**
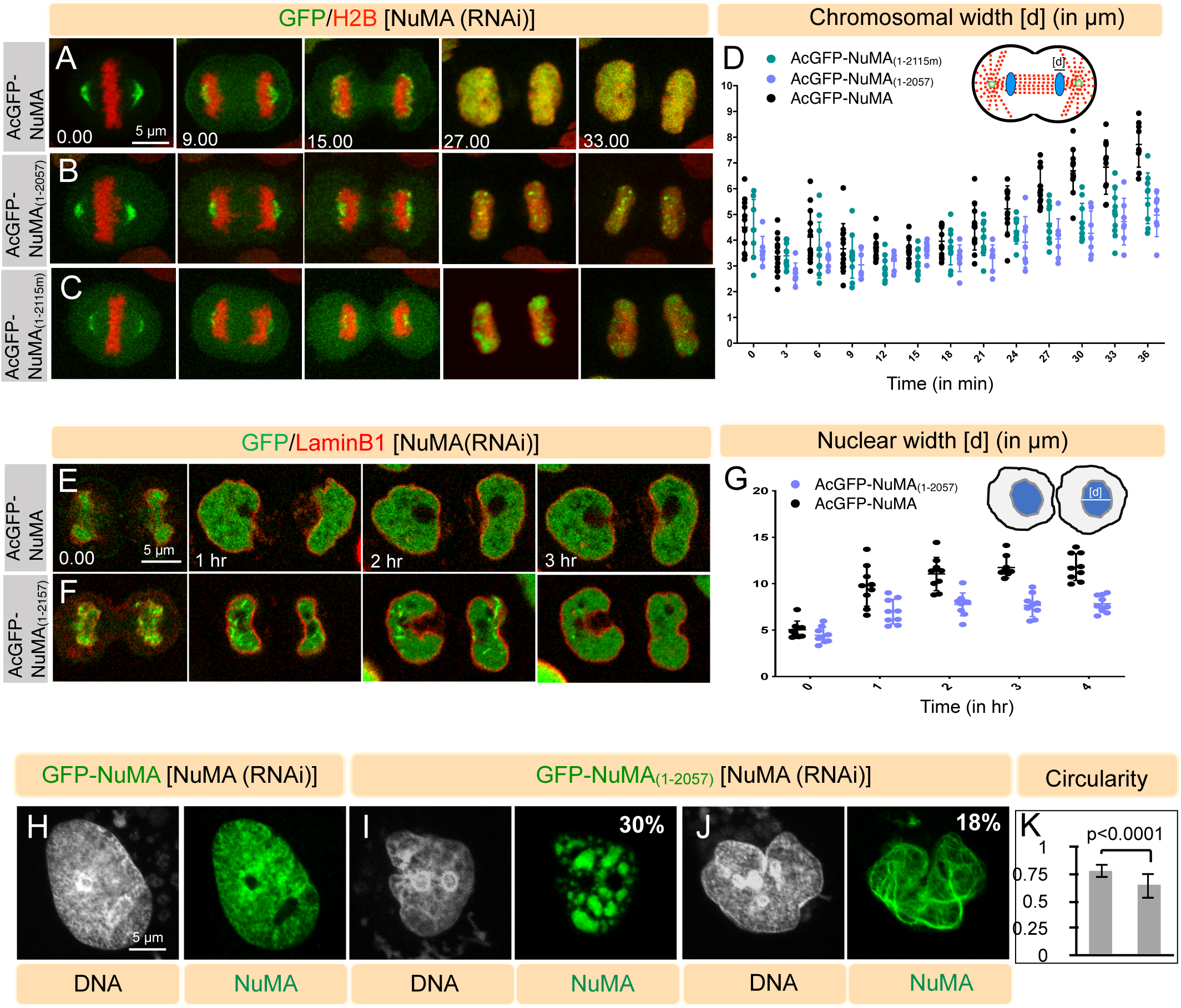
NuMA-DNA is vital for DNA decompaction and proper nuclear architecture. (A-C) Images from the time-lapse confocal microscopy of HeLa cells stably expressing mCherry-H2B and depleted of endogenous NuMA by RNAi using siRNAs sequences targeting 3’UTR of NuMA (see the depletion efficiency of siRNAs in Fig. 1B). These cells, as indicated, are transfected with AcGFP-NuMA (A), AcGFP-NuMA_(1-2057)_ (B), or AcGFP-NuMA_(1-2115m)_(C). The GFP signal is shown in green, and the mCherry signal is in red. Time is in minutes (min). Time ‘0’ min being the last frame of metaphase before the onset of chromosomes segregation (also see corresponding Supplementary Movies S4-S6). (D) Schematic representation for the measurement of the chromosomal width ([d] in µm) for the cells shown in A-C for 36 min and their quantification. Note significantly reduced chromosomal width in cells expressing AcGFP-NuMA_(1-2057)_ and AcGFP-NuMA_(1-2115m)_ when compared to AcGFP-NuMA from 27 min onwards (p<0.001 for t=27s, 30s, 33s, 36s; error bars: SD) (E, F) Images from the time-lapse confocal microscopy of HeLa Kyoto cells stably expressing mCherry-LaminB1 and depleted of endogenous NuMA by RNAi using siRNAs sequences targeting 3’UTR of NuMA. These cells, as indicated, are transfected with either AcGFP-NuMA (E) or AcGFP-NuMA_(1-2057)_ (F). The GFP signal is shown in green, the mCherry signal in red. Time is indicated in hours (hr). Time’ 0’ is the time when AcGFP-NuMA_(1-2115)_ or AcGFP-NuMA_(1-2057)_ enters the nucleus at the nuclear envelope reformation, which is similar for both constructs (data not shown). (G) Schematic representation for the measurement of the nuclear width ([d] in µm) for the cells shown in E and F on the left, and their quantification. Quantification was performed by measuring the maximum width of maximum intensity projected confocal images during the early G1 phase of the cell cycle. Please note the significant reduced nuclear width in cells expressing AcGFP-NuMA_(1-2057)_. (H-J) HeLa Kyoto cells in interphase are partly depleted of endogenous NuMA by RNAi using siRNAs sequences targeting 3’UTR of NuMA and transfected with AcGFP-NuMA (H) or AcGFP-NuMA_(1-2057)_ (I, J). Cells were stained for GFP (green), and DNA is visualized in gray. Note the cells that express AcGFP-NuMA_(1-2057)_ form puncta and fibrillar structure that are completely missing in from cells expressing wild-type form of NuMA (see also Supplementary Figure S5A-S5E). Also, see the impact of AcGFP-NuMA_(1-2057)_ expression on the nuclear shape. The percentage of cells showing puncta or fibrillar structure is indicated in the images. (K) Quantification of circularity (see materials and methods) of nuclei in cells expressing AcGFP-NuMA, or AcGFP-NuMA_(1-2057)_ (n=70 cells; error bars: SD).

A significant impact on chromatin decompaction in the early G1 observed above with NuMA_(1-2057)_ expression prompted us to examine the nuclear architecture of cells that weakly express AcGFP-NuMA_(1-2057)_ in NuMA (RNAi) background. Notably, we found a dramatic impact on the nuclear shape in cells expressing these constructs in comparison with the full-length wild-type protein (Fig. 4H-4K). The effect of NuMA_(1-2057)_ expression on interphase nuclei cannot be attributed to their role in mitosis, as the expression of these constructs entirely suppress the mitotic abnormalities seen upon endogenous NuMA depletion (Supplementary Fig. S1A and S1B). Interestingly, a significant number of cells expressing NuMA_(1-2057)_ that showed irregular nuclear shape were characterized by the presence of either puncta or fibrillar structure that were never seen in wild-type NuMA expressing cells (compare Fig. 4I, 4J with 4H). Also, the presence of these structures in cells expressing NuMA_(1-2057)_ was merely not due to over-expression, as comparable over-expression of the wild-type protein in cells did not lead to the formation of such structures (Supplementary Fig. S5A-S5E). Overall, these data suggest that NuMA-DNA interaction is vital for timely DNA decompaction during mitotic exit and proper nuclear shape in the interphase nuclei.

### DNA binding ability of NuMA is essential for preventing its higher-order assembly in interphase nuclei

To further characterize the origin of puncta or fibrillar structure in AcGFP-NuMA_(1-2057)_ expressing cells, we conducted live recording in HeLa cells that were stably expressing mCherry-H2B and transiently transfected with AcGFP-NuMA_(1-2057)_. Results obtained from this analysis revealed that within 9 hr of live-cell recording, [24%] of cells show soluble protein to puncta formation, [14%] of cells show soluble to fibrillar network, and small populations of cells [5%] form fibrillar assemblies from puncta (Fig. 5A-5C; Supplementary Movies S6-S10). Next, we characterized the biophysical properties of punctate and fibrillar structures using FRAP. In comparison with the homogeneously distributed NuMA_(1-2057)_ or NuMA_(1-2115m)_ protein that exchanges rapidly in FRAP as mentioned before (Figure 1), these higher-order assemblies of NuMA seen with NuMA_(1- 2057)_ or NuMA_(1-2115m)_ expression were significantly slow in their recovery profile (Supplementary Fig. S5F-S5M).

**Figure 5.**
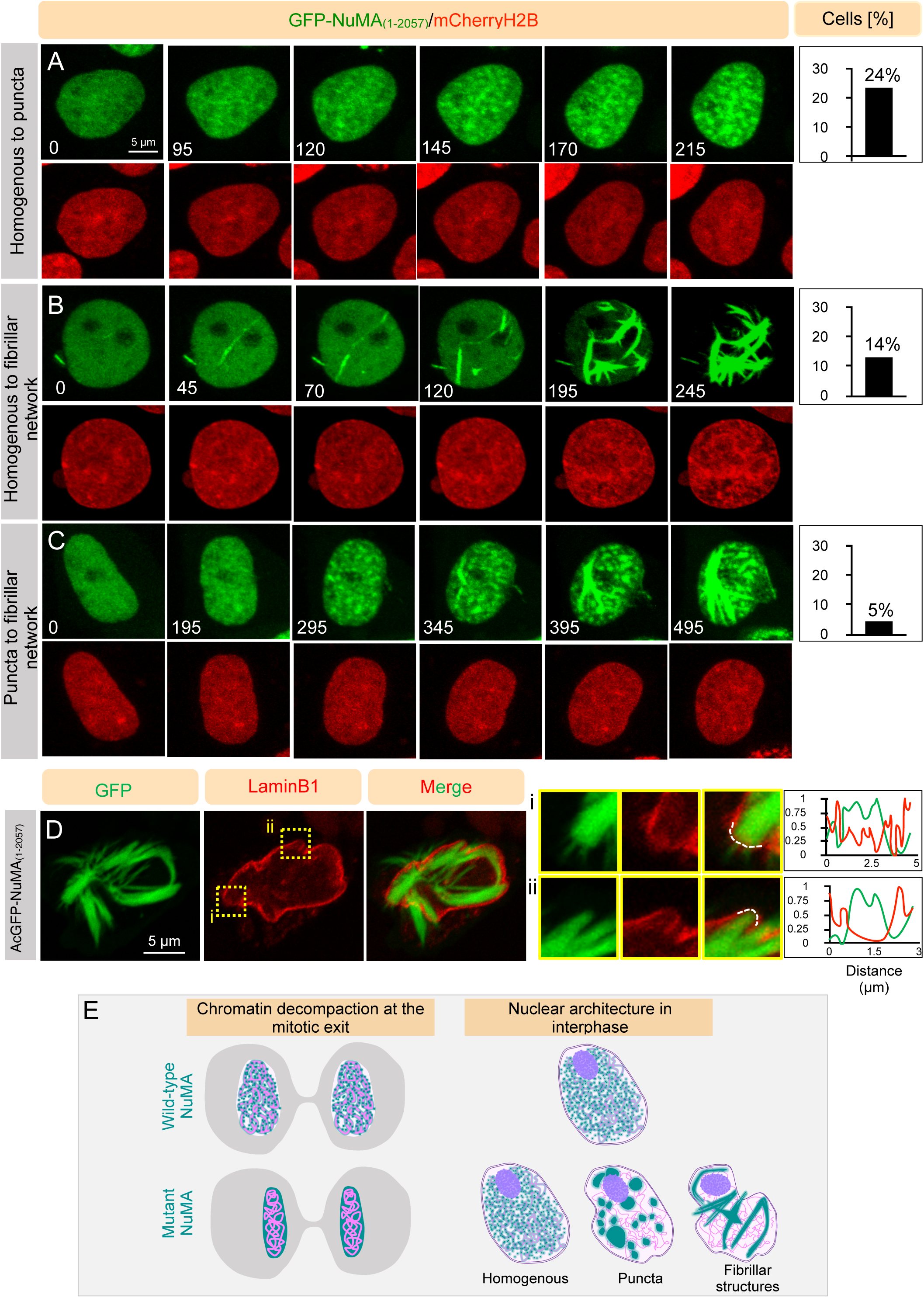
NuMA lacking the DNA binding potential assembles into higher-order structures in the nucleus. (A-C) Images from the long-term time-lapse recording of HeLa cells stably expressing mCherry-H2B and transfected with AcGFP-NuMA_(1-2057)_. The expression of AcGFP-NuMA_(1-2057)_ leads to higher-order assemblies within the nucleus. These assemblies categorized into three groups: homogenous to puncta formation (A), homogenous to the solid fibrillar network (B), or puncta to the solid fibrillar network (C). Quantification on the right represents the [%] of cells that are grouped into these categories while conducting 9 hr of live-imaging. (n=68 cells). Also, see corresponding Supplementary Movies S7-S10. (D) Images from the time-lapse recording of HeLa Kyoto cells stably expressing mCherry-LaminB1 and transfected with AcGFP-NuMA_(1-2057)_. The expression of AcGFP-NuMA_(1-2057)_ leads to solid fibrillar networks. Insets of the areas (i and ii) is shown on the right with the line-scan intensity of mCherry-LaminB1 and of AcGFP-NuMA_(1-2057)_ signal at the dashed while line covering a portion of the nuclear envelope. Note the decrease in the mCherry-LaminB1 intensity at those regions where AcGFP-NuMA_(1-2057)_-based fibrillar network are mechanically rupturing the nuclear envelope. (n>10 cells). (E) Model for the NuMA function during mitotic exit and in the interphase nuclei. In the control cells, wild-type NuMA interacts with chromatin during nuclear envelope reformation, and this allows chromatin to de-compact in telophase/early G1 phase of the cell cycle. However, in the absence of such an interaction, chromatin remains compact, and thus the width of the nucleus in the newly formed daughter cells remains significantly smaller. Also, such mutant NuMA exists in three different forms: homogenous, puncta, and solid fibrillar network. And, these higher-order assemblies, including puncta and solid fibrillar network mechanically deform the nuclear architecture.

This observation prompted us to test if these higher-order solid fibrillar structures based on NuMA_(1-2057)_ expression are responsible for nuclear deformation by mechanically pressing against the nuclear envelope. To this end, we analyzed AcGFP-NuMA_(1-2057)_ expressing cells that form the solid fibrillar structure in the HeLa cell line that is stably expressing mCherry-Lamin B1 and also depleted for the endogenous NuMA protein. Remarkably, NuMA-based solid fibrillar assemblies that form in the proximity of the nuclear envelope were capable of mechanically deforming the nuclear envelope. This feature was never detected in cells expressing AcGFP-tag wild-type NuMA protein (Fig. 5D, and data not shown). Overall, these sets of results suggest that the binding of NuMA to the DNA prevents higher order NuMA assemblies comprising of puncta and solid fibrillar structure, and these structures are fatal for the nuclear architecture.

Chromatin in the interphase nucleus is non-randomly distributed into defined regions called chromosomal territories (Andrew Fritz et al., 2016). During nuclear envelope reassembly (NER) at the mitotic exit, a cell ensures that these territories are maintained while the genetic material is undergoing decondensation. In the past few years, several proteins have been linked with the proper chromosomes decondensation at the mitotic exit, for instance, PP1 phosphatase, Aurora B kinase, and p97 AAA+ ATPase (Vagnarelli et al., 2011; Ramadan et al., 2007). However, despite these efforts, our knowledge about the nature of proteins and their spatiotemporal role in chromosome decompaction remains incomplete. Here, we identify the novel contribution of a well-established mitotic regulator NuMA in orchestrating chromosome decompaction during mitotic exit. We present evidence that NuMA directly associates the chromatin through the evolutionarily conserved arginine and lysine-rich sequences present in its C-terminus. In the absence of such an interaction, chromatin decompaction is perturbed (Fig. 5E). How NuMA promote correct decondensation of the chromatin at the mitotic exit? Previous work has shown that purified NuMA protein can assembles into multiarms oligomers (Gueth-Hallonet et al., 1998; Harborth et al., 1999).

And, overexpression of NuMA can lead to the formation of a three-dimensional quasi-hexagonal lattice in mammalian cells (Harborth et al., 1999). Based on these observations, we propose that the interaction of NuMA to the chromatin is necessary to keep the chromatin in a proper organization at the mitotic exit. In the absence of such association, chromatin remains tightly packaged (Fig. 5E). It would be interesting to determine the functional significance of this phenotype for gene regulation in the early G1 phase of the cell cycle. Since the formation of NuMA dimer by the coiled-coil interaction is the first key step for the oligomerization assembly, we assume that the NuMA lacking the coiled-coil domain would also be deficient in proper chromatin decondensation at the mitotic exit. Therefore, dissecting the function of the coiled-coil in chromosomes decompaction would be an exciting step to discover the hidden feature of this crucial molecule further.

We demonstrated that Cdk1 activity helps in releasing NuMA from the chromatin in the late-prophase. This outcome could be because of the direct phosphorylation of NuMA by Cdk1, or an indirect consequence of Cdk1 phosphorylation on some other yet unknown protein. Because NuMA directly associates with the DNA *in vitro* and C-terminus fragments of NuMA comprising the last 355 aa or smaller are efficient in binding to the chromosomes at all the mitotic stages, we favor the model that direct phosphorylation by Cdk1 dissociates NuMA from the chromosomes in mitosis.

Unfortunately, mutation of all the Cdk1-phosphorylated residues mapped in the several phosphoproteomics data one by one did not lead to the identification of Cdk1-regulated residues/s that is responsible for releasing NuMA from the chromatin. We envisage that this could be due to redundancy between more than one Cdk1-regulated residue in regulating NuMA-chromatin interaction.

A population of DNA-binding deficient mutant of NuMA that is localized as a homogenous protein in the nucleoplasm showed a faster [t_1/2_] recovery rate in comparison with full-length protein. However, nuclei in these cells remain spherical and do not reveal any nuclear shape phenotype (Fig. 1). Therefore, it would be interesting to mechanically challenge cells that expresses homogenously distributed DNA-binding deficient mutant of NuMA to characterize its function in providing mechanical stability to the interphase nucleus. Remarkably, a significant number of nuclei expressing this mutant protein also showed higher-order assembly of mutated NuMA into puncta and solid fibrillar networks. And, the nuclei that carry these structures are improper in their architecture (Fig. 5E). Notably, we uncovered that the solid fibrillar networks in these nuclei could mechanically deform the nucleus by compressing onto the nuclear membrane (Fig. 5E). Our data further rule out the possibility that the change in nuclear architecture is due to abnormal mitosis, as the expression of these mutated fragments fully rescues mitotic abnormality observed upon endogenous protein depletion.

Nonetheless, establishing a strategy for knocking out or knocking down NuMA only in the interphase nuclei would be essential to address the function of endogenous protein for nuclear shape.

Since errors in maintaining the proper nuclear shape and gene expression is associated with pathological disorders such as cancer and aging (Zink et al., 2004; Dahl et al., 2008), we believe that our effort in identifying the NuMA’s role in orchestrating correct nuclear architecture will be instrumental in broadening our knowledge in this arena.

## Materials and Methods

### Cell culture, plasmid, and siRNAs transfection, and stable cell line generation

HeLa cells, hTERT-RPE1 cells, and HeLa Kyoto cells stably expressing mCherry-H2B, mCherry-LaminB1, AcGFP-NuMA were cultured in high-glucose DMEM with GlutaMAX (CC3004; Genetix) supplemented with 10% fetal calf serum (FCS) in a humidified 5% CO2 incubator at 37°C. For plasmid transfections, cells were seeded at 80% confluency in imaging dish (0030740017; Eppendorf) or on coverslips in 6-well plates. 4 µg of plasmid DNA suspended in 400 µl of serum-free DMEM was incubated for 5 min, followed by the addition of 6 µl of Lipofectamine 2000 (11668019; Life Technologies), was mixed and incubated for 15 min. This mixture was then added to the cells. And the cells were fixed or imaged 30-36 hr post-transfection.

For siRNA experiment, 6 or 9 µl of 20 µM siRNA and 4 µl of Lipofectamine RNAi MAX (13778150, Invitrogen) were suspended in 100 µl of water (W4502; Sigma) and were incubated for 5 min in parallel, then mixed and incubated for another 15 min. This mixture was added to the 2.5 ml medium per well containing around 100,000 cells. Cells were then grown for 60-72 hr before fixation or live-imaging.

For the generation of stable cells expressing AcGFP-NuMA, AcGFP-H2B, and mCherry-LaminB1 in HeLa Kyoto, cells were cultured in 10 cm dish at 80% confluency. These cells were then transfected with 6 µg of pIRES-AcGFP-FLAG-NuMA, pIRES-H2B-AcGFP or pIRES-mCherry-LaminB1 plasmid using 12µl of Lipofectamine. After 36 hr, 400ng/µl puromycin media was added for the selection. Isolated colonies were cultured, and clones were confirmed by immunostaining and immunoblot analysis.

### Plasmids and siRNAs

All NuMA clones were amplified from a previously existing plasmid as a template with appropriate PCR primer pairs. NuMA full-length and NuMA_(1-2057)_ was cloned into pIRES-AcGFP-FLAG plasmid (a gift from Mark Petronczki) using Age1 and EcoR1 site. For cloning NuMA_(1-2115m)_, Kpn2I site was introduced in pIRES-AcGFP-NuMA plasmid and custom made double-stranded DNA (from Macrogen, Inc.) where all the arginine and lysine residues from 2058-2115 are converted to alanine was cloned using Kpn2I and EcoR1 sites. pIRES-H2B-AcGFP Plasmid was cloned by replacing mCherry fragment from Addgene plasmid 21044 with amplified AcGFP using Age1 and Not1 site. pIRES-NLS-AcGFP plasmid was cloned by incorporating SV-40 NLS sequence in the forward primer used for amplifying AcGFP. All the smaller C-terminus fragments of NuMA was cloned in pcDNA3-GFP vector (Merdes et al., 2000) using Xba1 and Not1 site. All Cdk1 phosphorylated residues (Threonine or Serine) in NuMA were mutated using the megaprimer approach. For recombinant protein expression in *E. coli*, NuMA_(2058-2115)_, NuMA_(1877-2115)_ and NuMA_(1877-2057)_ were cloned in pET30a Plasmid with a hexa-histidine tag at N-terminus using Nco1 and EcoR1 sites. Bacteria histone-like HU protein was generously provided by V. Nagaraja (MCB, IISc).

Double-stranded siRNA oligonucleotides used were 5’- CAGUACCAGUGAGUGGCCCCACCUG-3’ (NuMA 3’UTR siRNA; Eurogentec) and 5’-CACCGUGUGUCUAAGCAAA-3’ (RCC1 siRNA; Eurogentec).

### Drug-mediated inhibition of mitotic kinases

hTERT-RPE1 cells were synchronized in early prophase by double thymidine block. Briefly, the cells were treated with 2 mM thymidine (T1895; Sigma-Aldrich) for 17 hr, released for 8 hr followed by another round of thymidine treatment. Cells were treated with DMSO for control, 20 µM or 100 µM of Cdk1 inhibitor RO-3306 (S7747; Selleckchem) for 10 min, 250 nM or 1.25 µM of Aurora A inhibitor MLN-8054 (S1100; Selleckchem) for 1 hr, 300 nM or 1.5 µM of Plk1 inhibitor BI-2536 (S1109; Selleckchem) for 30 min before fixation. Following treatment with the inhibitors, cells were fixed with cold methanol and immunostained with antibodies against NuMA (sc-48773; Santa Cruz) and γ-tubulin (GTU88; Sigma-Aldrich).

### Time-lapse imaging and FRAP analysis

Time-lapse microscopy was conducted on Olympus FV 3000 confocal laser scanning microscope using a 40X NA 1.3 oil (Olympus Corporation, Japan) using an imaging dish (0030740017; Eppendorf) at 5% CO2, 37°C, 90% humidity maintained by Tokai Hit STR Stage Top incubator with Touch panel Controller. For mitotic cells, images were acquired at the interval of either 2 min or 3 min with 9-11 optical sections (3 μm a part). For the interphase cells, images were captured every 5 min with optical sectioning of 1 μm.

FRAP experiments were performed for a specific region (1.75 µm^2^) of the nucleus of HeLa cells stably expressing AcGFP-NuMA, AcGFP-H2B or cells that are transiently transfected with AcGFP-NuMA_(1-2057)_, AcGFP-NuMA_(1-2115m)_ or NLS-AcGFP with a 40X objective. 40% of the 488-nm laser was utilized to bleach the region of interest, and images within the same focal plane were acquired for every 5 s for the entire duration of 50 cycles to monitor fluorescence recovery. Due to faster recovery in cells that are expressing AcGFP-NuMA_(1-2057)_, AcGFP-NuMA_(1-2115m)_ or NLS-AcGFP images were acquired every 2 s. To assess the fluorescence loss due to photobleaching, fluorescence from a region separated from the bleached region was simultaneously recorded. The intensity value in the bleached area was measured, corrected for the background, and the curves were then normalized using the following equation:

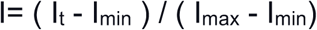

where I represents the normalized intensity, I_t_ represents the intensity at a time-point, I_min_ is the minimum intensity (at the time of bleaching), and I_max_ is the maximum intensity (pre bleaching intensity). For the calculation of half-time of recovery [t_1/2_] or mobile fraction, the bleaching due to imaging was considered, and the values were quantified by fitting to 1st order exponential equation using Origin software (https://www.originlab.com/origin).

### Nuclear fractionation

To obtain nuclear matrix or chromosomal fraction, we utilized a method as described in Abad *et al*, 2007. In brief, HeLa Kyoto cells washed with PBS-protease inhibitors (PI) (Merck: Cat no 539134) and were collected at 450g at 4°C for 5 min and were suspended in 1 ml of Buffer A (10 mM HEPES, pH 7.4, 1 mM EGTA, 2 mM MgCl2, 250 mM sucrose, and PI). After that, 1 ml of Buffer B (1 mM HEPES, pH 7.4, containing PI) was added and was incubated on ice for 30 min. Separation of nuclei from the cytoplasm was performed using Dounce homogenizer, and this was confirmed under an epifluorescence microscope using Hoechst 33342 (B2261; Sigma-Aldrich). The nuclear pellet was collected at 3200g at 4°C for 15 min and was suspended in buffer-X (10 mM HEPES, pH 7.9, 10 mM KCl, 1.5 mM MgCl2, 0.34 M sucrose, 10% (vol/vol) glycerol, 1 mM dithiothreitol, PI) with Triton X-100 0.1% (vol/vol), and was incubated on ice for 8 min, and pellet down at 1300g at 4°C for 5 min. The pellet was washed again with buffer Nuclei was lysed in Buffer Y (3 mM EDTA, 0.2 mM EGTA, 1mM dithiothreitol, and PI) on ice for 30 min. The pellet formed at 1650g at 4°C for 5 min was rewashed with Buffer The pellet was subjected to 0.1 µl Mnase (EN0181; Fermentas) in 100 µl of MNase Buffer (10 mM Tris, pH 8.8, 10 mM KCl, and 1mM CaCl2) at 37 °C for 15 min. The reaction was stopped using 1mM EGTA. The nuclease sensitive, Chromatin Fraction (Chr.), and the resistant matrix fraction (Matrix) was separated by at 1650g at 4°C for 5min. The MF was washed with 100 µl of MNase buffer. CF and MF suspended in Laemmli buffer and denatured at 99 °C for 10 min, and then utilized for immunoblotting.

### Super-resolution imaging

Super-resolution imaging was conducted on Olympus spinning disk super-resolution confocal microscope (IXplore SpinSR) using a 100X 1.45 NA Objective. Images were acquired by capturing 13 Z-sections, 0.23 µm apart. The images were processed with Olympus Super Resolution (OSR) software.

### Electrophoretic Mobility Shift Assay (EMSA)

Recombinant protein HU, NuMA_(2058-2115)_, NuMA_(1877-2115)_, NuMA_(1877-2057)_, or NuMA-Nter(1-705) was incubated with 400ng of PUC18 Plasmid in 1X-TAE (89mM tris, 89mM acetic acid, 1mM EDTA pH 8.4) buffer at 27°C for 30 min. The protein-DNA complexes were resolved in 4% acrylamide gel in 1X-TAE buffer. The gel was stained with ethidium bromide and visualized under UV light.

### Indirect immunofluorescence and immunoblotting

For immunofluorescence, cells were fixed with chilled methanol at -20°C for 10 min and washed in PBST (PBS containing 0.05% Triton X-100). Cells were blocked in 1% BSA (RM3159; HiMedia) for 1 hr, followed by incubation with primary antibody for 4 hours at room temperature. After three washes of 5 min each with PBST, cells were incubated with secondary antibody for 1 hr. Cells were then given three washes with PBST and stained with 1 µg/ml Hoechst 33342 (B2261; Sigma-Aldrich) for 5 minutes. Following three washes with PBST, the coverslips were mounted using Fluoromount (SouthernBiotech, 0100-01). The primary antibodies used were 1:1000 mouse anti-GFP (2955S; Cell signalling), 1:200 rabbit anti-NuMA (sc-48773; Santa-Cruz), 1:200 mouse anti-RCC1 (sc-376049; Santa-Cruz), 1:1000 mouse anti-γ-tubulin (GTU88; Sigma-Aldrich), 1:200 rabbit anti-Ect2 (07-1364; Merck). Secondary antibodies used were 1:500 Alexa flour 488 goat anti-mouse (A11001; Invitrogen), 1:500 Alexa flour 488 goat anti-rabbit (A11008; Invitrogen), 1:500 Alexa flour 568 goat anti-mouse (A11004; Invitrogen), and 1:500 Alexaflour 568 goat anti-rabbit (A11011; Invitrogen). Confocal images were acquired on Olympus FV 3000 confocal laser scanning microscope using 60X NA 1.4 oil objective. All the images are processed in ImageJ.

For immunoblotting, HeLa Kyoto cells or HeLa Kyoto cells transfected with NuMA siRNAs or HeLa Kyoto cells stably expressing AcGFP-NuMA, synchronized in prometaphase with 100 nM Nocodazole (M1404; Sigma-Aldrich) for 20 hr. Cells were lysed in lysis buffer (50 mM Tris, pH-7.4, 2 mM EDTA, 2 mM EGTA, 25 mM Sodium fluoride, 0.1 mM sodium orthovanadate, 0.1 mM PMSF, 0.2% Triton-X100, 0.3% NP-40, 100 nM Okadaic acid, and complete EDTA-free protease inhibitor) for 2 hr on ice and after a spin of 14000 rpm, cell supernatant was denatured at 99°C in 2X SDS–PAGE buffer and analyzed by SDS–PAGE followed by immunoblotting. For immunoblotting, 1:1000 rabbit anti-NuMA (sc-48773; Santa Cruz), 1:5000 mouse anti-actin (sc-58673; Santa Cruz), 1:1000 of mouse anti-LaminB1 (sc-6216; Santa-Cruz), and rabbit anti-RNA polymerase A (sc-899; Santa-Cruz) antibodies were used.

### Quantifications and statistical analysis

All quantifications were performed in ImageJ. Quantification of GFP chromosomal intensity was determined by calculating the ratio of mean chromosomal intensity and mean cytoplasmic intensity (of a rectangular region of interest of area 1.69 µm^2^) and correcting for the background signal.

For chromosomal width measurements, a line was drawn on the longest width of chromatin on the maximum intensity projection images, and the length of the line was measured.

The circularity of the nucleus was calculated using the freehand tool to manually select the outline of the nucleus, and circularity was calculated using the formula 4π(area/perimeter^2^).

Spindle pole enrichment of NuMA was determined by calculating the ratio of mean spindle pole intensity and mean cytoplasmic intensity after correcting for background signal, as described in Sana et al., 2018.

Midzone Ect2 intensity was measured using a rectangular region of interest of 1.09 µm^2^ and was corrected for background signal.

Whole-cell GFP intensity was measured using the freehand tool in ImageJ to select the outline of the nucleus. To rule out the difference in nuclear area of individual cells, the intensity was divided by the total area used for quantification to obtain intensity/µm^2^.

To calculate the significance of the differences between two mean values, unpaired *t* test was used. *p*-value was considered to be significant if *p*≤ 0.05 using GraphPad Prism 8.

## Acknowledgements

We are grateful to Sophie Dumont and Andrea Serra-Marques (UCSF) for sharing with us their unpublished data, and for the fruitful discussions. We thank Andreas Merdes (CBI, Toulouse), Arnaud Echard (Institut Pasteur), Mark Petronczki (Boehringer Ingelheim, Vienna), V. Nagaraja (IISc, Bangalore) and Anthony Hyman (MPI-CBG, Dresden) for providing us plasmids, recombinant protein, and cell lines. We are grateful to Phong Tran, Raj Ladher, Sveta Chakrabarti, Fernando R Balestra, and Sukriti Kapoor for providing us critical comments on the manuscript. We thank Ganesh Kadasoor (Olympus Corporation) for taking super-resolution images on the Olympus IXplore SpinSR microscope system, and Sukriti Kapoor for making a working model for us. We thank DST-FIST, UGC Centre for the Advanced Study, DBT-IISc Partnership Program, and IISc for the infrastructure support. This work is supported by the grants from the Wellcome Trust DBT-India Alliance [grant number: IA/I/15/2/502077] to SK. SK is a Wellcome Trust DBT-India Alliance Intermediate Fellow.

## Notes

### Competing Interest Statement

The authors have declared no competing interest.

## References

Abad, P.C., J. Lewis, I.S. Mian, D.W. Knowles, J. Sturgis, S. Badve, J. Xie, and S.A. Lelievre. 2007. NuMA influences higher order chromatin organization in human mammary epithelium. Mol Biol Cell. 18:348–361.

Antonin, W., and H. Neumann. 2016. Chromosome condensation and decondensation during mitosis. Curr Opin Cell Biol. 40:15–22.

Aureille, J., V. Buffiere-Ribot, B.E. Harvey, C. Boyault, L. Pernet, T. Andersen, G. Bacola, M. Balland, S. Fraboulet, L. Van Landeghem, and C. Guilluy. 2019. Nuclear envelope deformation controls cell cycle progression in response to mechanical force. EMBO Rep. 20:e48084.

Batty, P., and D.W. Gerlich. 2019. Mitotic Chromosome Mechanics: How Cells Segregate Their Genome. Trends Cell Biol. 29:717–726.

Cleveland, D.W. 1995. NuMA: a protein involved in nuclear structure, spindle assembly, and nuclear re-formation. Trends Cell Biol. 5:60–64.

Compton, D.A., and D.W. Cleveland. 1993. NuMA is required for the proper completion of mitosis. J Cell Biol. 120:947–957.

Compton, D.A., I. Szilak, and D.W. Cleveland. 1992. Primary structure of NuMA, an intranuclear protein that defines a novel pathway for segregation of proteins at mitosis. J Cell Biol. 116:1395–1408.

Cremer, T., and M. Cremer. 2010. Chromosome territories. Cold Spring Harb Perspect Biol. 2:a003889.

Cuijpers, S.A.G., and A.C.O. Vertegaal. 2018. Guiding Mitotic Progression by Crosstalk between Post-translational Modifications. Trends Biochem Sci. 43:251–268.

Dahl, K.N., A.J. Ribeiro, and J. Lammerding. 2008. Nuclear shape, mechanics, and mechanotransduction. Circ Res. 102:1307–1318.

Das, A.T., M.E. Luderus, and W.H. Lamers. 1993. Identification and analysis of a matrix-attachment region 5’ of the rat glutamate-dehydrogenase-encoding gene. Eur J Biochem. 215:777–785.

Dephoure, N., C. Zhou, J. Villen, S.A. Beausoleil, C.E. Bakalarski, S.J. Elledge, and S.P. Gygi. 2008. A quantitative atlas of mitotic phosphorylation. Proc Natl Acad Sci U S A. 105:10762–10767.

Du, Q., and I.G. Macara. 2004. Mammalian Pins is a conformational switch that links NuMA to heterotrimeric G proteins. Cell. 119:503–516.

Endo, A., A. Moyori, A. Kobayashi, and R.W. Wong. 2013. Nuclear mitotic apparatus protein, NuMA, modulates p53-mediated transcription in cancer cells. Cell Death Dis. 4:e713.

Finn, E.H., G. Pegoraro, H.B. Brandao, A.L. Valton, M.E. Oomen, J. Dekker, L. Mirny, and T. Misteli. 2019. Extensive Heterogeneity and Intrinsic Variation in Spatial Genome Organization. Cell. 176:1502-1515.e1510.

Friedl, P., K. Wolf, and J. Lammerding. 2011. Nuclear mechanics during cell migration. Curr Opin Cell Biol. 23:55–64.

Fritz, A.J., A.R. Barutcu, L. Martin-Buley, A.J. van Wijnen, S.K. Zaidi, A.N. Imbalzano, J.B. Lian, J.L. Stein, and G.S. Stein. 2016. Chromosomes at Work: Organization of Chromosome Territories in the Interphase Nucleus. J Cell Biochem. 117:9–19.

Gueth-Hallonet, C., J. Wang, J. Harborth, K. Weber, and M. Osborn. 1998. Induction of a regular nuclear lattice by overexpression of NuMA. Exp Cell Res. 243:434–452.

Harborth, J., J. Wang, C. Gueth-Hallonet, K. Weber, and M. Osborn. 1999. Self assembly of NuMA: multiarm oligomers as structural units of a nuclear lattice. Embo j. 18:1689–1700.

Harborth, J., K. Weber, and M. Osborn. 2000. GAS41, a highly conserved protein in eukaryotic nuclei, binds to NuMA. J Biol Chem. 275:31979–31985.

Hoar, K., A. Chakravarty, C. Rabino, D. Wysong, D. Bowman, N. Roy, and J.A. Ecsedy. 2007. MLN8054, a small-molecule inhibitor of Aurora A, causes spindle pole and chromosome congression defects leading to aneuploidy. Mol Cell Biol. 27:4513–4525.

Houtsmuller, A.B. 2005. Fluorescence recovery after photobleaching: application to nuclear proteins. Adv Biochem Eng Biotechnol. 95:177–199.

Hubner, M.R., M.A. Eckersley-Maslin, and D.L. Spector. 2013. Chromatin organization and transcriptional regulation. Curr Opin Genet Dev. 23:89–95.

Hueschen, C.L., V. Galstyan, M. Amouzgar, R. Phillips, and S. Dumont. 2019. Microtubule End-Clustering Maintains a Steady-State Spindle Shape. Curr Biol. 29:700-708.e705.

Kallajoki, M., J. Harborth, K. Weber, and M. Osborn. 1993. Microinjection of a monoclonal antibody against SPN antigen, now identified by peptide sequences as the NuMA protein, induces micronuclei in PtK2 cells. J Cell Sci. 104 (Pt 1):139–150.

Kallajoki, M., K. Weber, and M. Osborn. 1991. A 210 kDa nuclear matrix protein is a functional part of the mitotic spindle; a microinjection study using SPN monoclonal antibodies. Embo j. 10:3351–3362.

Kallajoki, M., K. Weber, and M. Osborn. 1992. Ability to organize microtubules in taxol-treated mitotic PtK2 cells goes with the SPN antigen and not with the centrosome. J Cell Sci. 102 (Pt 1):91–102.

Kiyomitsu, T., and I.M. Cheeseman. 2012. Chromosome- and spindle-pole-derived signals generate an intrinsic code for spindle position and orientation. Nat Cell Biol. 14:311–317.

Kotak, S., K. Afshar, C. Busso, and P. Gonczy. 2016. Aurora A kinase regulates proper spindle positioning in C. elegans and in human cells. J Cell Sci. 129:3015–3025.

Kotak, S., C. Busso, and P. Gonczy. 2012. Cortical dynein is critical for proper spindle positioning in human cells. J Cell Biol. 199:97–110.

Kotak, S., C. Busso, and P. Gonczy. 2013. NuMA phosphorylation by CDK1 couples mitotic progression with cortical dynein function. Embo j. 32:2517–2529.

Kotak, S., C. Busso, and P. Gonczy. 2014. NuMA interacts with phosphoinositides and links the mitotic spindle with the plasma membrane. Embo j. 33:1815–1830.

Lammerding, J., P.C. Schulze, T. Takahashi, S. Kozlov, T. Sullivan, R.D. Kamm, C.L. Stewart, and R.T. Lee. 2004. Lamin A/C deficiency causes defective nuclear mechanics and mechanotransduction. J Clin Invest. 113:370–378.

Lindqvist, A., V. Rodriguez-Bravo, and R.H. Medema. 2009. The decision to enter mitosis: feedback and redundancy in the mitotic entry network. J Cell Biol. 185:193–202.

Lydersen, B.K., and D.E. Pettijohn. 1980. Human-specific nuclear protein that associates with the polar region of the mitotic apparatus: distribution in a human/hamster hybrid cell. Cell. 22:489–499.

Merdes, A., and D.W. Cleveland. 1998. The role of NuMA in the interphase nucleus. J Cell Sci. 111 (Pt 1):71–79.

Merdes, A., R. Heald, K. Samejima, W.C. Earnshaw, and D.W. Cleveland. 2000. Formation of spindle poles by dynein/dynactin-dependent transport of NuMA. J Cell Biol. 149:851–862.

Merdes, A., K. Ramyar, J.D. Vechio, and D.W. Cleveland. 1996. A complex of NuMA and cytoplasmic dynein is essential for mitotic spindle assembly. Cell. 87:447–458.

Mirny, L.A., M. Imakaev, and N. Abdennur. 2019. Two major mechanisms of chromosome organization. Curr Opin Cell Biol. 58:142–152.

Misteli, T. 2007. Beyond the sequence: cellular organization of genome function. Cell. 128:787–800.

Mueller, F., D. Mazza, T.J. Stasevich, and J.G. McNally. 2010. FRAP and kinetic modeling in the analysis of nuclear protein dynamics: what do we really know? Curr Opin Cell Biol. 22:403–411.

Nagano, T., Y. Lubling, C. Varnai, C. Dudley, W. Leung, Y. Baran, N. Mendelson Cohen, S. Wingett, P. Fraser, and A. Tanay. 2017. Cell-cycle dynamics of chromosomal organization at single-cell resolution. Nature. 547:61–67.

Nishimoto, T., E. Eilen, and C. Basilico. 1978. Premature of chromosome condensation in a ts DNA-mutant of BHK cells. Cell. 15:475–483.

Nishitani, H., M. Ohtsubo, K. Yamashita, H. Iida, J. Pines, H. Yasudo, Y. Shibata, T. Hunter, and T. Nishimoto. 1991. Loss of RCC1, a nuclear DNA-binding protein, uncouples the completion of DNA replication from the activation of cdc2 protein kinase and mitosis. Embo j. 10:1555–1564.

Nozaki, T., R. Imai, M. Tanbo, R. Nagashima, S. Tamura, T. Tani, Y. Joti, M. Tomita, K. Hibino, M.T. Kanemaki, K.S. Wendt, Y. Okada, T. Nagai, and K. Maeshima. 2017. Dynamic Organization of Chromatin Domains Revealed by Super-Resolution Live-Cell Imaging. Mol Cell. 67:282-293.e287.

Petronczki, M., M. Glotzer, N. Kraut, and J.M. Peters. 2007. Polo-like kinase 1 triggers the initiation of cytokinesis in human cells by promoting recruitment of the RhoGEF Ect2 to the central spindle. Dev Cell. 12:713–725.

Petrone, A., M.E. Adamo, C. Cheng, and A.N. Kettenbach. 2016. Identification of Candidate Cyclin-dependent kinase 1 (Cdk1) Substrates in Mitosis by Quantitative Phosphoproteomics. Mol Cell Proteomics. 15:2448–2461.

Price, C.M., and D.E. Pettijohn. 1986. Redistribution of the nuclear mitotic apparatus protein (NuMA) during mitosis and nuclear assembly. Properties of purified NuMA protein. Exp Cell Res. 166:295–311.

Radulescu, A.E., and D.W. Cleveland. 2010. NuMA after 30 years: the matrix revisited. Trends Cell Biol. 20:214–222.

Ramadan, K., R. Bruderer, F.M. Spiga, O. Popp, T. Baur, M. Gotta, and H.H. Meyer. 2007. Cdc48/p97 promotes reformation of the nucleus by extracting the kinase Aurora B from chromatin. Nature. 450:1258–1262.

Rogers, S., R.A. McCloy, B.L. Parker, R. Chaudhuri, V. Gayevskiy, N.J. Hoffman, D.N. Watkins, R.J. Daly, D.E. James, and A. Burgess. 2015. Dataset from the global phosphoproteomic mapping of early mitotic exit in human cells. Data Brief. 5:45–52.

Sana, S., R. Keshri, A. Rajeevan, S. Kapoor, and S. Kotak. 2018. Plk1 regulates spindle orientation by phosphorylating NuMA in human cells. Life Sci Alliance. 1:e201800223.

Sekiya, T., U.M. Muthurajan, K. Luger, A.V. Tulin, and K.S. Zaret. 2009. Nucleosome-binding affinity as a primary determinant of the nuclear mobility of the pioneer transcription factor FoxA. Genes Dev. 23:804–809.

Seldin, L., N.D. Poulson, H.P. Foote, and T. Lechler. 2013. NuMA localization, stability, and function in spindle orientation involve 4.1 and Cdk1 interactions. Mol Biol Cell. 24:3651–3662.

Simon, D.N., and K.L. Wilson. 2011. The nucleoskeleton as a genome-associated dynamic ‘network of networks’. Nat Rev Mol Cell Biol. 12:695–708.

Smith, E.R., Y. Meng, R. Moore, J.D. Tse, A.G. Xu, and X.X. Xu. 2017. Nuclear envelope structural proteins facilitate nuclear shape changes accompanying embryonic differentiation and fidelity of gene expression. BMC Cell Biol. 18:8.

Steegmaier, M., M. Hoffmann, A. Baum, P. Lenart, M. Petronczki, M. Krssak, U. Gurtler, P. Garin-Chesa, S. Lieb, J. Quant, M. Grauert, G.R. Adolf, N. Kraut, J.M. Peters, and W.J. Rettig. 2007. BI 2536, a potent and selective inhibitor of polo-like kinase 1, inhibits tumor growth in vivo. Curr Biol. 17:316–322.

Tabellini, G., M. Riccio, G. Baldini, R. Bareggi, A.M. Billi, V. Grill, P. Narducci, and A.M. Martelli. 2001. Further considerations on the intranuclear distribution of HMGI/Y proteins. Ital J Anat Embryol. 106:251–260.

Tang, T.K., C.J. Tang, Y.L. Chen, and C.W. Wu. 1993. Nuclear proteins of the bovine esophageal epithelium. II. The NuMA gene gives rise to multiple mRNAs and gene products reactive with monoclonal antibody W1. J Cell Sci. 104 (Pt 2):249–260.

Vagnarelli, P., S. Ribeiro, L. Sennels, L. Sanchez-Pulido, F. de Lima Alves, T. Verheyen, D.A. Kelly, C.P. Ponting, J. Rappsilber, and W.C. Earnshaw. 2011. Repo-Man coordinates chromosomal reorganization with nuclear envelope reassembly during mitotic exit. Dev Cell. 21:328–342.

van Steensel, B., and A.S. Belmont. 2017. Lamina-Associated Domains: Links with Chromosome Architecture, Heterochromatin, and Gene Repression. Cell. 169:780–791.

Vassilev, L.T., C. Tovar, S. Chen, D. Knezevic, X. Zhao, H. Sun, D.C. Heimbrook, and L. Chen. 2006. Selective small-molecule inhibitor reveals critical mitotic functions of human CDK1. Proc Natl Acad Sci U S A. 103:10660–10665.

Webster, M., K.L. Witkin, and O. Cohen-Fix. 2009. Sizing up the nucleus: nuclear shape, size and nuclear-envelope assembly. J Cell Sci. 122:1477–1486.

Woodard, G.E., N.N. Huang, H. Cho, T. Miki, G.G. Tall, and J.H. Kehrl. 2010. Ric-8A and Gi alpha recruit LGN, NuMA, and dynein to the cell cortex to help orient the mitotic spindle. Mol Cell Biol. 30:3519–3530.

Yang, C.H., E.J. Lambie, and M. Snyder. 1992. NuMA: an unusually long coiled-coil related protein in the mammalian nucleus. J Cell Biol. 116:1303–1317.

Yang, C.H., and M. Snyder. 1992. The nuclear-mitotic apparatus protein is important in the establishment and maintenance of the bipolar mitotic spindle apparatus. Mol Biol Cell. 3:1259–1267.

Zeng, C., D. He, and B.R. Brinkley. 1994. Localization of NuMA protein isoforms in the nuclear matrix of mammalian cells. Cell Motil Cytoskeleton. 29:167–176.

Zheng, Z., Q. Wan, G. Meixiong, and Q. Du. 2014. Cell cycle-regulated membrane binding of NuMA contributes to efficient anaphase chromosome separation. Mol Biol Cell. 25:606–619.

Zink, D., A.H. Fischer, and J.A. Nickerson. 2004. Nuclear structure in cancer cells. Nat Rev Cancer. 4:677–687.

