## Supplementary text and figures for "NuMA interaction with chromatin is vital for proper nuclear architecture in human cells"

Supplementary Information

Supplemental Figure Legends

Supplemental Figure 1

NuMA interacts with the chromatin in the interphase nucleus

(A, B) Characterization of chromosomal instability during metaphase (A), and chromosomal bridges in anaphase (B) in HeLa Kyoto cells which are transfected with scrambled siRNAs (Control), siRNAs against NuMA 3'-UTR for 72 hr which are either left untreated or transiently transfected with AcGFP-NuMA, AcGFP-NuMA$_{(1-2057)}$ or AcGFP-NuMA$_{(1-2115m)}$. Note the complete rescue of these phenotypes in the cells expressing AcGFP-NuMA, AcGFP-NuMA$_{(1-2057)}$, or AcGFP-NuMA$_{(1-2115m)}$. (n>40 cells and the experiment is done twice; error bars: SD).

(C-F) FRAP analysis of HeLa Kyoto cells that are stably expressing AcGFP-H2B (C, E) or transiently transfected with AcGFP-NLS (D, F). The GFP signal is shown in green, and the time is indicated in seconds (s). The unbleached and bleached region of the cell is shown by yellow and white circles, respectively. The GFP recovery profile of the bleached area corrected for photobleaching is plotted for 200s for AcGFP-H2B and 14s for AcGFP-NLS. The half-time of recovery [$t_{1/2}$] of these cells is shown in the main Fig. 1I. (n=9 cells for AcGFP-NLS and n=7 for AcGFP-H2B expressing cells Error bars: SD).

(G) Immunoblot analysis of protein extracts prepared from the isolated nuclei from HeLa Kyoto cells, which are later fractionated into MNase sensitive chromatin fraction, and a nuclear matrix fraction (see Materials and Methods for details). These fractions were probed with antibodies against NuMA, LaminB1 (a marker for insoluble matrix fraction), and RNA Pol II subunit a (that mainly associates with chromatin fraction). The molecular mass is indicated in kilodaltons (kDa). Note that a significant amount of NuMA is present in the chromatin fraction.

Supplemental Figure 2

RanGEF RCC1 is not critical for NuMA enrichment at chromatin in prophase

(A-C) hTERT-RPE1 cells in prophase transfected with scrambled siRNAs (Control) (A), siRNAs against RCC1 (B), or NuMA siRNAs (C). Cells were fixed after 72 hr and stained for RCC1 (green) and NuMA (red); DNA is shown in blue. In this and other figures, a white line on the image indicates the region used for the line scan plot that is shown on the right. Note that NuMA and RCC1 localization on chromatin is independent of each other.

(D) HeLa Kyoto cell in prophase transfected with GFP-NuMA$_{(1411-2115)}$. Cells were fixed after 36 hr and stained for GFP (green). The line scan plot is shown on the right.

Supplemental Figure 3

Cdk1/cyclinB1 activity is critical for releasing NuMA from chromatin at NEBD

(A-C) HeLa cells in metaphase were treated either with DMSO (Control) (A) or Aurora A inhibitor MLN-8054 (250 nM for 1 hr) (B) before fixation. These cells were stained for NuMA (red) and γ-tubulin (green). DNA is shown in blue. Quantification on the right shows spindle pole enrichment of NuMA in cells treated with MLN-8054 (C).
(D-F) HeLa cells in anaphase that were either treated with DMSO (Control) (D) or Plk1 inhibitor BI-2536 (300 nM for 30 min) (E) before fixation. These cells were stained for Ect2 (green). DNA is shown in blue. Quantification on the right shows central spindle enrichment for Ect2.
(G-J) hTERT-RPE1 cells synchronized in late prophase, as indicated in for figure panel 2A, are treated either with DMSO control (G), or 5X concentration of the inhibitor MLN-8054 (H), BI-2536 (I) or RO-3306 (J) that is reported for Fig. 2D-2F. After fixation, these cells are stained for NuMA (red) and γ-tubulin (green). Linescan plot is shown on the right. Note the non-retention of NuMA on chromatin in prometaphase cells even when the cells were treated with 5X concentration of Aurora A and Plk1 inhibitors, suggesting that Aurora A and Plk1 are not involved in releasing NuMA from chromatin at NEBD. The percentage of cells showing chromosomal retention of NuMA is indicated on the image. (n>20 cells in each condition and experiments were repeated twice).
(K) Threonine or Serine residues that are reported to get phosphorylated by Cdk1 in the various phosphoproteomics data set. These residues were mutated to Alanine to analyze NuMA localization at the metaphase stage of HeLa cells. None of these

mutations resulted in the retention of NuMA at the chromosomes in late mitosis, suggesting they may act redundantly.

Supplemental Figure 4

Amino acid sequences 2058-2115 of NuMA are vital for chromatin localization

(A-E) Prophase or prometaphase stages as indicated from the time-lapse recording of HeLa cells stably expressing mCherry-H2B and transiently transfected with GFP-NuMA$_{(1411-2115)}$ (A), GFP-NuMA$_{(1700-2115)}$ (B), GFP-NuMA$_{(1760-2115)}$ (C), GFP-NuMA$_{(1991-2115)}$ (D) or GFP-NuMA$_{(2058-2115)}$ (E). Time is indicated in minutes with t=0 corresponding to late prophase before NEBD. Note that in prophase, expression of all the fragments is localized to chromatin, however, in prometaphase upon NEBD, only GFP-NuMA$_{(1760-2115)}$, GFP-NuMA$_{(1991-2115)}$ and GFP-NuMA$_{(2058-2115)}$ are retained on chromatin. Also, see the corresponding Supplementary Movie S1 for cell expressing GFP-NuMA$_{(1411-2115)}$; S2 for cell expressing GFP-NuMA$_{(2058-2115)}$, and S3 for cells expressing GFP-NuMA$_{(1411-2057)}$.

(F, G) Images from the time-lapse recording of HeLa cells stably expressing mCherry-H2B and transiently transfected with either GFP-NuMA$_{(1760-2115)}$ (F) or GFP-NuMA$_{(1760-2057)}$ (G). Note that as shown here in F or the main Fig. 3D GFP-NuMA$_{(1760-2115)}$ localizes at the chromosomes all the time in mitosis. However, expression of GFP-NuMA$_{(1760-2057)}$ construct that lacks the last 58 aa is unable to localize at the chromosome in metaphase and early anaphase and only localized to the chromosome mass in late anaphase once the nuclear membrane is formed. Time is indicated in minutes.

(H) Quantification of the ratio of chromosomal to cytoplasmic intensity of GFP-NuMA$_{(1760\text{-}2115)}$ and GFP-NuMA$_{(1760\text{-}2057)}$ over time. Note that GFP-NuMA$_{(1760\text{-}2057)}$ is not localized on chromatin in the early stages of mitosis compared to GFP-NuMA$_{(1760\text{-}2115)}$. $p < 0.0001$ for all time points. ($n=10$ cells; Error bars: SD).

Supplemental Figure 5

<u>Higher-order assemblies formation with the non-DNA binding mutants of NuMA</u>

(A-E) HeLa Kyoto cell in interphase that are transfected with AcGFP-NuMA (A) or AcGFP-NuMA$_{(1\text{-}2057)}$. AcGFP-NuMA$_{(1\text{-}2057)}$ expression leads to the formation of higher-order assemblies as puncta (B), small fibrillar network (C), or large fibrillar network (D). Cells were fixed after 36 hr and stained for GFP (green). GFP intensity quantification on the right in (E) per $\mu$m2 in cells expressing either wild-type full-length AcGFP-NuMA construct or AcGFP-NuMA$_{(1\text{-}2057)}$ that forms either puncta, small or large fibrillar network. More than 10 cells were used for quantification in each condition. Note the formation of puncta or fibrillar network is not dependent on the amount of GFP-expression.

(F-M) FRAP analysis of puncta and fibrillar structure in HeLa cells that are stably expressing mCherry-H2B and are depleted of endogenous NuMA by using siRNAs against 3'UTR. These cells were transiently transfected either with AcGFP-NuMA$_{(1\text{-}2057)}$ (F, G, J, L) or AcGFP-NuMA$_{(1\text{-}2115m)}$ (H, I, K, M). The unbleached and bleached region is shown by yellow and white circles, respectively. Time is indicated in seconds. The recovery profile of the bleached area corrected for photobleaching is plotted for 80s (J, K) (Error bar: SD). Note highly delayed recovery profile of the fibrillar network in contrast to the puncta. Also, note that the mobile fraction of AcGFP-NuMA$_{(1\text{-}2057)}$ and AcGFP-NuMA$_{(1\text{-}2115m)}$ is significantly reduced for the puncta and fibrillar structure in comparison with the homogenous form that is remarkably mobile (L, and M) (n>9 cells; Error bar: SEM).

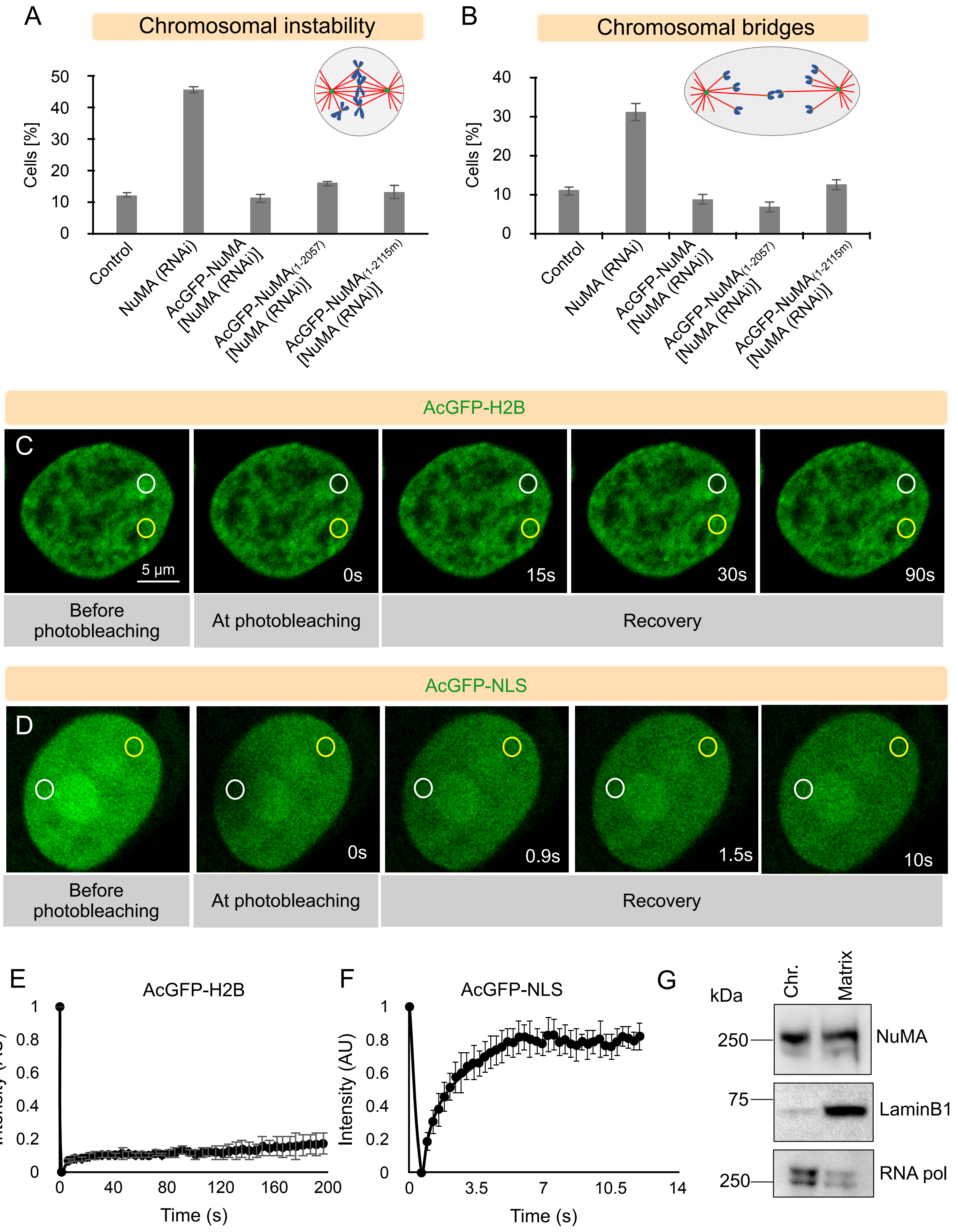


Supplementary Figure 1

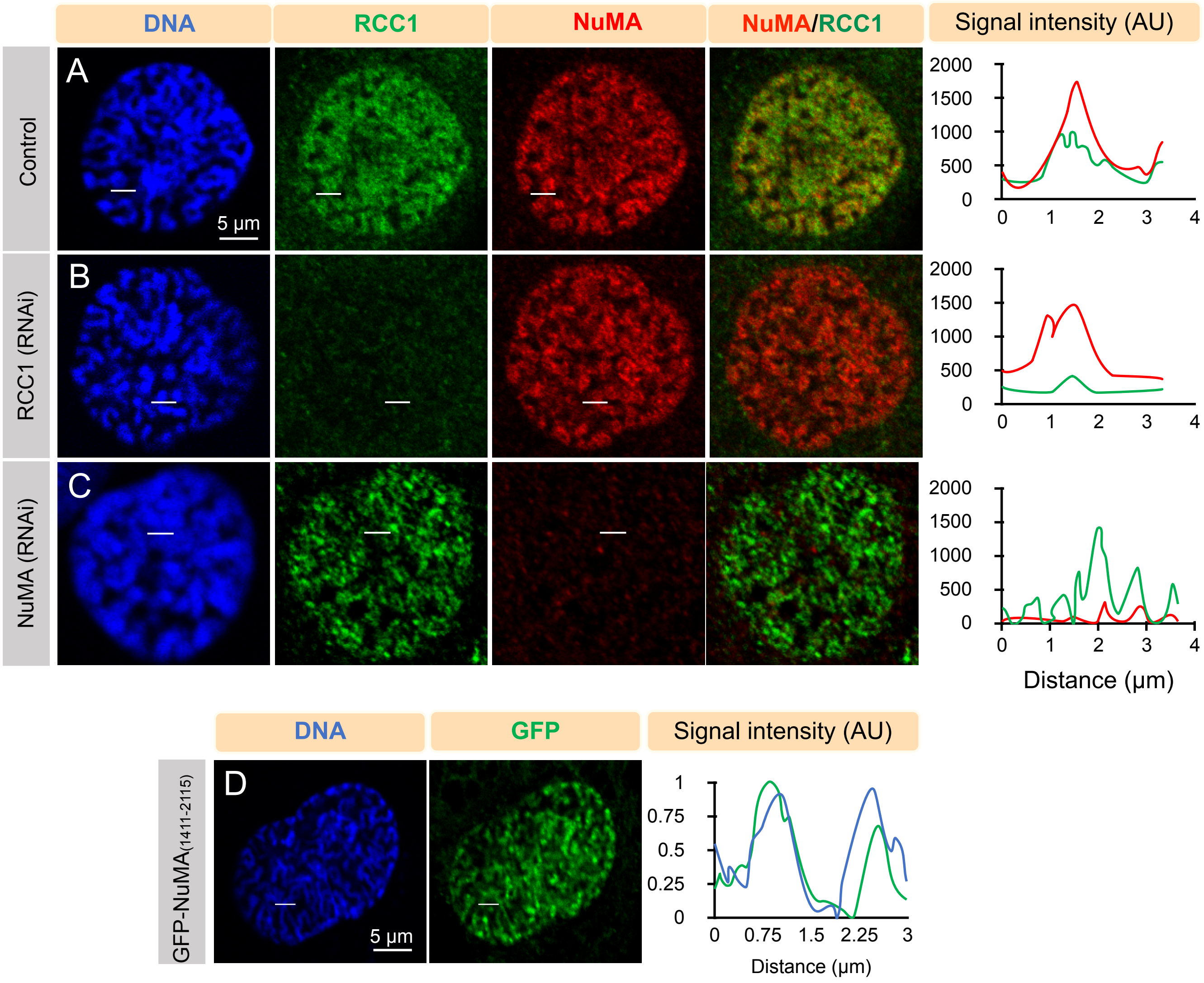


Supplementary Figure 2

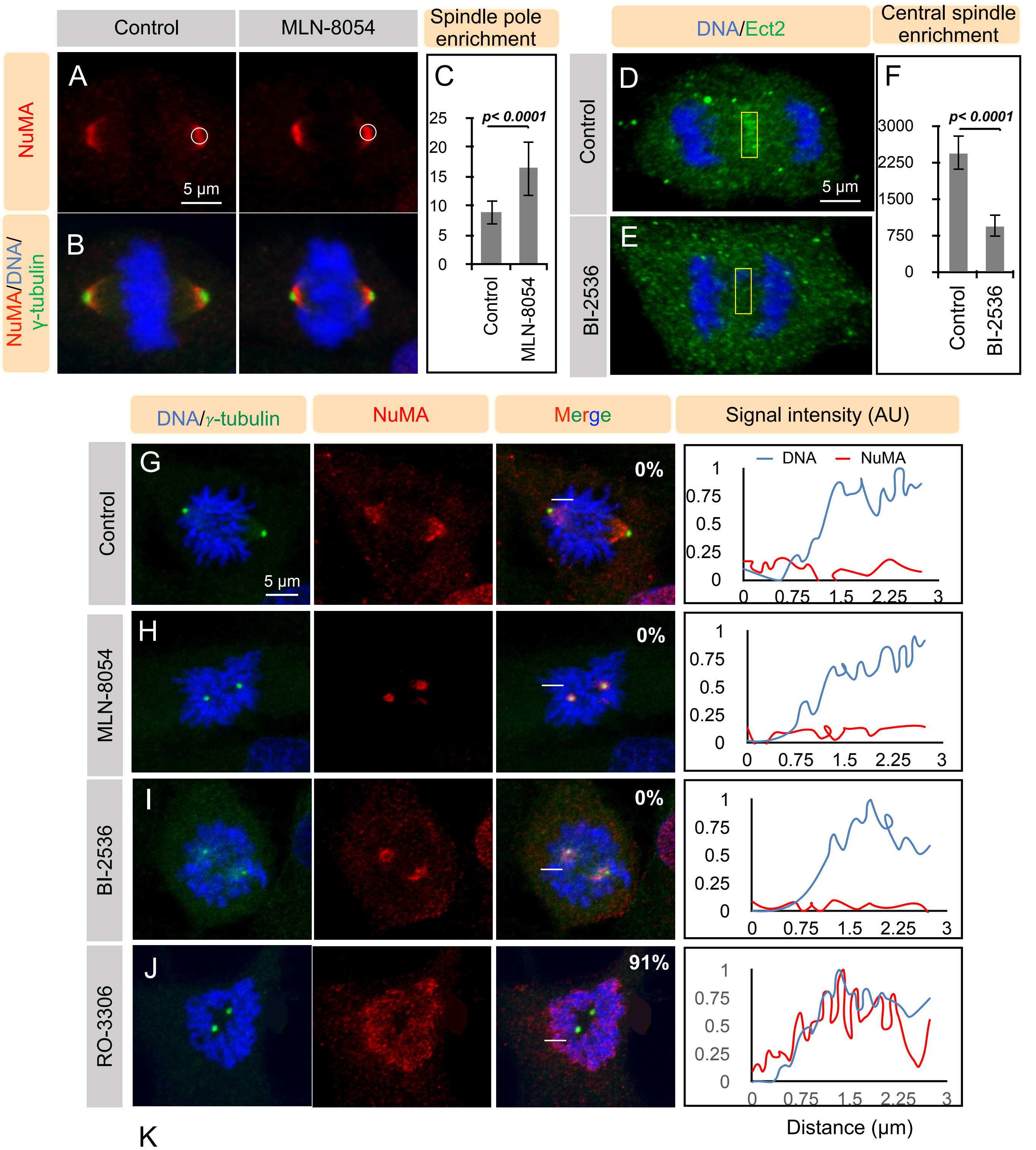


K

| Sl. No. | Cdk1 site on NuMA | Sequence | Reference | Metaphase chromosomal localization |
|---|---|---|---|---|
| 1 | T211 | GDILQTPQFQM | Rogers *et al*, 2015 | No |
| 2 | S271 | KQAASPLEPK | Rogers *et al*, 2015 | No |
| 3 | T1733 | CEEGTPLSIT | Dephoure *et al*, 2008 | No |
| 4 | S1757 | GEPASPISQR | Dephoure *et al*, 2008 | No |
| 5 | T1776 | ESLYFTPIPA | Rogers *et al*, 2015 | No |
| 6 | S1830 | VEEPDSANSSF | Petrone *et al*, 2016 | No |
| 7 | S1837 | ANSSFYSTRSAP | Petrone *et al*, 2016 | No |
| 8 | T2000 | QGPGTPESK | Rogers *et al*, 2015; Petrone *et al*, 2016 | No |
| 9 | S2047 | QADRRQSMAF | Rogers *et al*, 2015; Petrone *et al*, 2016 | No |

Supplementary Figure 3

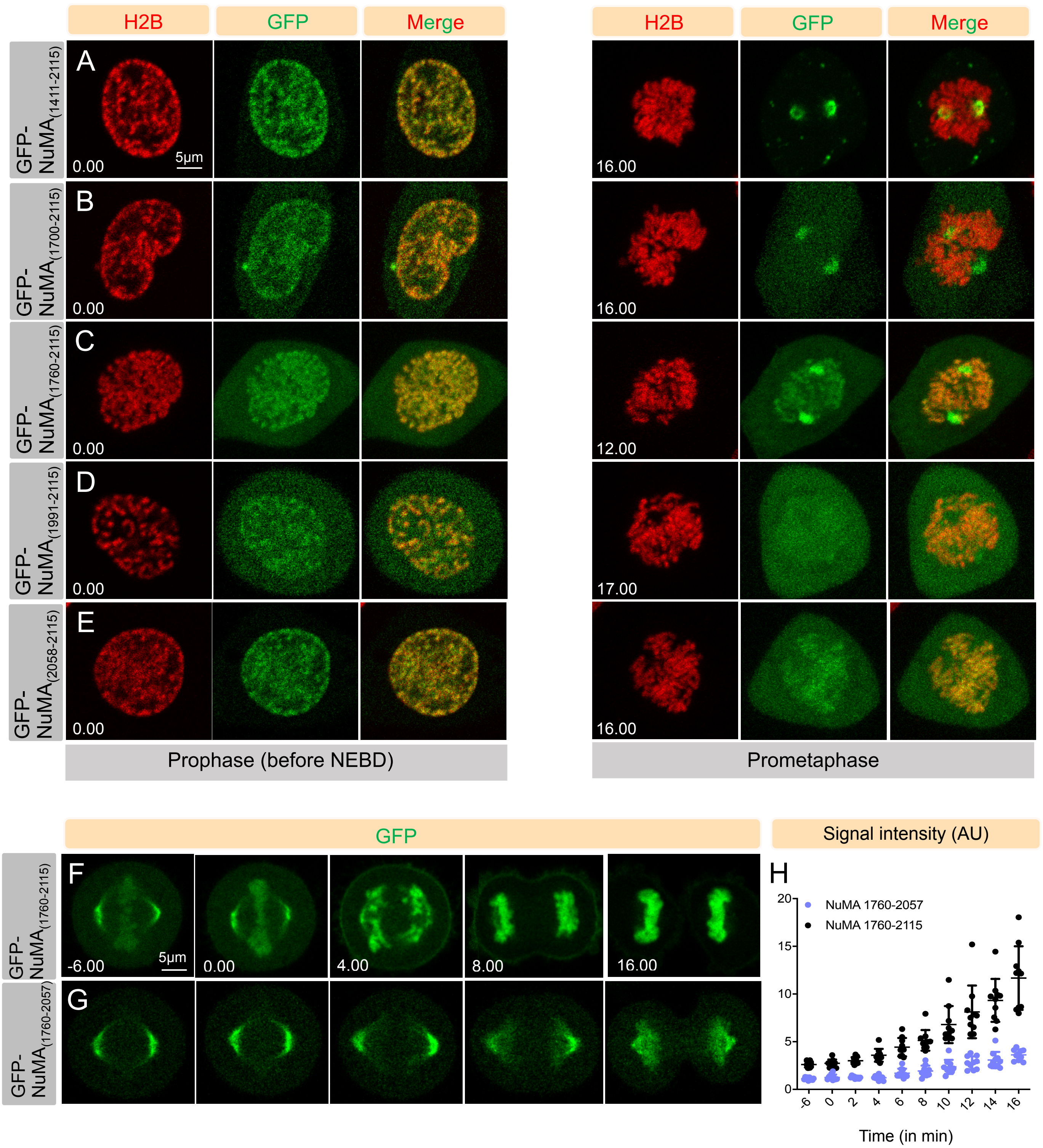


Supplementary Figure 4

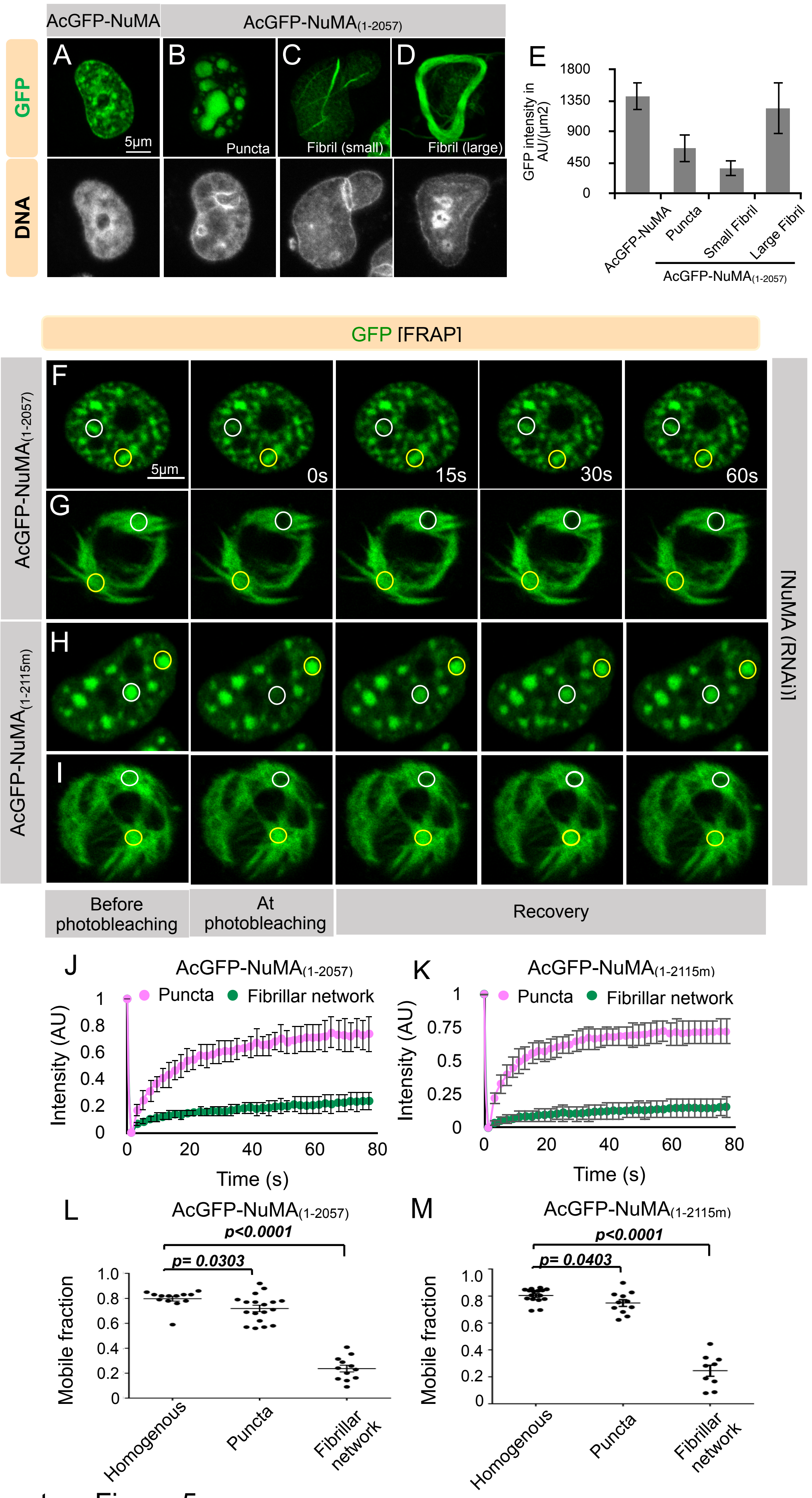


Supplementary Figure 5
